# Parametric control of flexible timing through low-dimensional neural manifolds

**DOI:** 10.1101/2021.11.08.467806

**Authors:** Manuel Beiran, Nicolas Meirhaeghe, Hansem Sohn, Mehrdad Jazayeri, Srdjan Ostojic

## Abstract

Biological brains possess an unparalleled ability to adapt behavioral responses to changing stimuli and environments. How neural processes enable this capacity is a fundamental open question. Previous works have identified two candidate mechanisms: a low-dimensional organization of neural activity and a modulation by contextual inputs. We hypothesized that combining the two might facilitate generalization and adaptation in complex tasks. We tested this hypothesis in the framework of flexible timing tasks where dynamics play a key role. Examining trained recurrent neural networks, we found that confining the dynamics to a low-dimensional subspace allowed tonic inputs to parametrically control the overall input-output transform, enabling generalization to novel inputs and adaptation to changing conditions. Reverse-engineering and theoretical analyses demonstrated that this parametric control relies on a mechanism where tonic inputs modulate the dynamics along non-linear manifolds in activity space while preserving their geometry. Comparisons with data from behaving monkeys confirmed the behavioral and neural signatures of this mechanism.

## 1 Introduction

Humans and animals can readily adapt and generalize their behavioral responses to new environmental conditions (Markman, 1989; Körding and Wolpert, 2004; Courville et al., 2006; Lake et al., 2015; Monosov, 2020). This capacity has been particularly challenging to realize in artificial systems (Lake et al., 2017; Sinz et al., 2019; Saxe et al., 2021), raising the question of what biological mechanism might enable it. Two mechanistic components have been put forward. A first proposed component is a low-dimensional organization of neural activity, a ubiquitous experimental observation across a large variety of behavioral paradigms (Gao et al., 2015; Gallego et al., 2017; Saxena and Cunningham, 2019; Jazayeri and Ostojic, 2021). Indeed, theoretical analyses have argued that, while high dimensionality promotes stimulus discrimination, low dimensionality instead facilitates generalization to previously unseen stimuli and conditions (DiCarlo and Cox, 2007; DiCarlo et al., 2012; Rigotti et al., 2013; Fusi et al., 2016; Chung et al., 2018; Cayco-Gajic and Silver, 2019; Bernardi et al., 2020; Nogueira et al., 2021). A second suggested component are tonic inputs that can help cortical network modulate their outputs to stimuli that appear in distinct behavioral context (Rigotti et al., 2010; Mante et al., 2013; Saez et al., 2015; Remington et al., 2018a; Cueva et al., 2020; Bernardi et al., 2020; Badre et al., 2021; Flesch et al., 2022; Naumann et al., 2022), and thereby generalize their responses to new circumstances. These two mechanisms have so far been investigated separately in different sets of simplified tasks, but they are not mutually exclusive and could in principle have complementary functions. Whether and how an interplay between low-dimensional dynamics and tonic inputs might enable generalization and adaptation in more complex cognitive tasks remains an open question.

Here we address this question within the framework of timing tasks that demand flexible control of the temporal dynamics of neural activity (Buonomano and Maass, 2009; Mello et al., 2015; Gouvêea et al., 2015; Merchant and Averbeck, 2017; Wang et al., 2018; Remington et al., 2018a,b; Gámez et al., 2019; Sohn et al., 2019; Egger et al., 2019; Cueva et al., 2020; Bi and Zhou, 2020; Monteiro et al., 2021; Meirhaeghe et al., 2021). Previous studies have demonstrated that a low-dimensional organization is a prominent feature of neural activity recorded during such tasks (Wang et al., 2018; Remington et al., 2018a,b; Sohn et al., 2019; Egger et al., 2019). Several of those studies moreover found that tonic inputs which provide contextual information allow networks to flexibly adjust their output to identical stimuli (Remington et al., 2018b; Sohn et al., 2019). However, so far, the computational roles of low-dimensional dynamics and contextual inputs were only probed within the range on which the network was trained. Going beyond previous works, we instead hypothesized that combining the two mechanisms by pairing contextual input signals with low-dimensional dynamics might facilitate generalization to novel inputs and adaptation to changing environments.

We investigate this hypothesis using a multidisciplinary approach spanning network modeling, theory and analyses of neural and behavioral data in non-human primates. We analyzed recurrent neural networks (RNNs) trained on a set of flexible timing tasks, where the goal was to produce a time interval depending on various types of inputs (Wang et al., 2018; Remington et al., 2018b; Sohn et al., 2019). To investigate the role of the dimensionality of neural trajectories, we exploited the class of low-rank neural networks, in which the dimensionality of dynamics is directly constrained by the rank of the recurrent connectivity matrix (Mastrogiuseppe and Ostojic, 2018; Schuessler et al., 2020a; Beiran et al., 2021; Dubreuil et al., 2022). We then compared the performance, generalization capabilities, and neural dynamics of these low-rank RNNs with networks trained in absence of any constraint on the connectivity and the dimensionality of dynamics. We similarly contrasted networks with and without tonic inputs.

We first show that low-rank RNNs, unlike unconstrained networks, are able to generalize to novel stimuli well beyond their training range, but only when paired with tonic inputs that provide contextual information. Specifically, we found that in low-rank networks, tonic inputs parametrically control the input-output transform and its extrapolation. Reverse-engineering the trained networks revealed that tonic inputs endowed the system with generalization by modulating the dynamics along low-dimensional non-linear manifolds while leaving their geometry invariant. Population analyses of neural data recorded in behaving monkeys performing a time-interval reproduction task (Sohn et al., 2019; Meirhaeghe et al., 2021) confirmed the key signatures of this geometry-preserving mechanism. In a second step, we show how the identified mechanism allows networks to quickly adapt their outputs to changing conditions by adjusting the tonic input based on the temporal statistics. Direct confrontation of newly collected behavioral and neural data with low-rank networks supported the idea that contextual inputs are key to adaptation. Altogether, our work brings forth evidence for computational benefits of controlling low-dimensional activity through tonic inputs, and provides a mechanistic framework for understanding the role of resulting neural manifolds in flexible timing tasks.

## 2 Results

### 2.1 Generalization in flexible timing tasks

We trained recurrent neural networks (RNNs) on a series of flexible timing tasks, in which networks received various contextual and/or timing inputs and had to produce a time interval that was a function of those inputs (Fig. 1 A, B). In all tasks, the network output was a weighted sum of the activity of its units (see Methods). This output was required to produce a temporal ramp following an externally provided ‘Set’ signal. The output time interval corresponded to the time between the ‘Set’ signal and the time at which the network output crossed a fixed threshold (Wang et al., 2018; Remington et al., 2018b; Sohn et al., 2019). Inputs to the network determined the target interval, and therefore the desired slope of the output ramp, but the nature of inputs depended on the task (Fig. 1 B). Our goal was to study which properties of RNNs favour generalization, i.e., extend their outputs to inputs not seen during training. We therefore examined the full input-output transform performed by the trained networks (Fig. 1 C), and specifically contrasted standard RNNs, which we subsequently refer to as “unconstrained RNNs”, with low-rank RNNs in which the connectivity was constrained to minimize the embedding dimensionality of the collective dynamics (Fig. 1 D, Dubreuil et al. (2022)).

**Figure 1:**
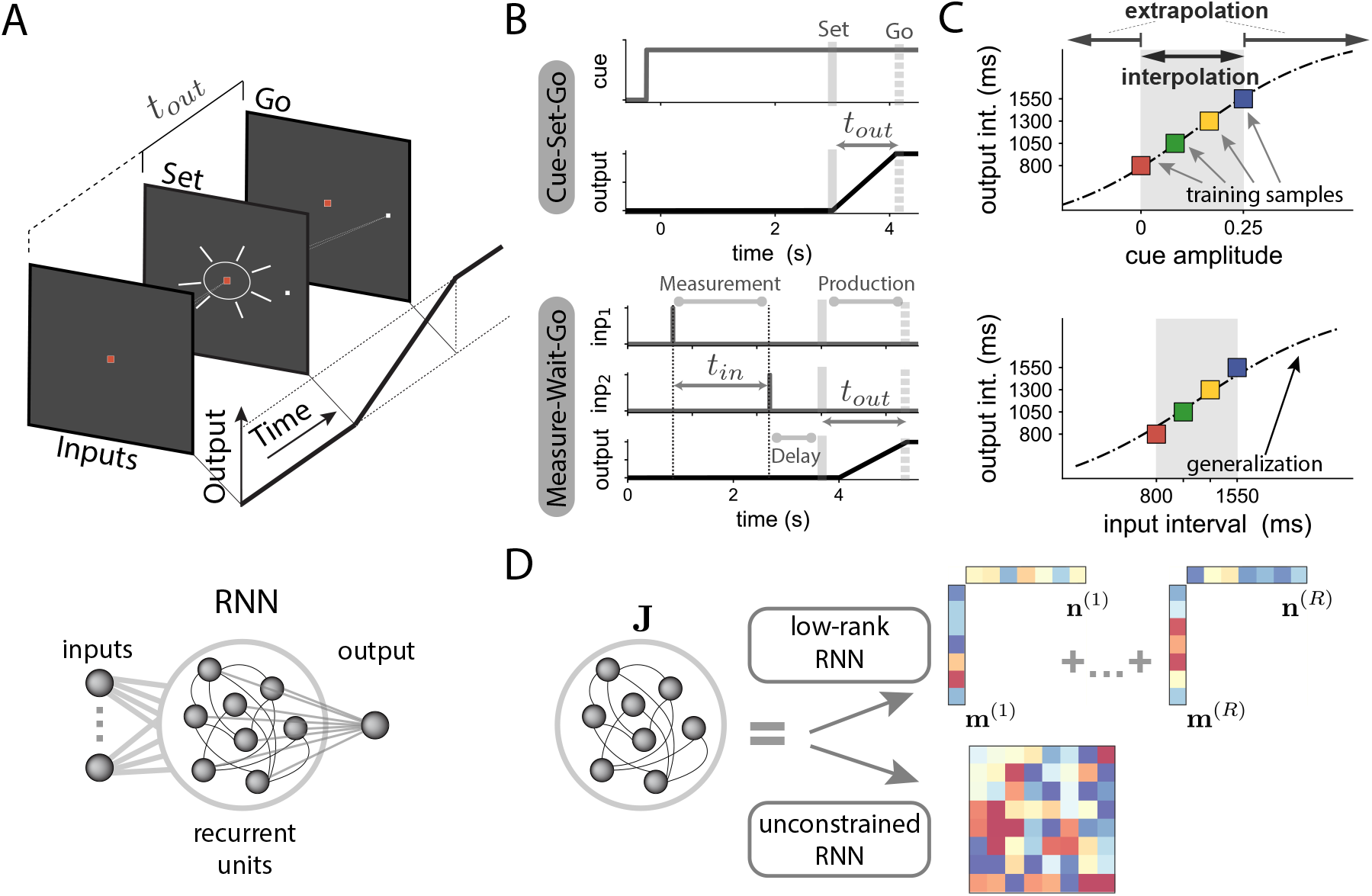
Investigating generalization in flexible timing tasks using low-rank recurrent neural networks (RNNs). **A** Flexible timing tasks. Top: On each trial, a series of timing inputs that depend on each task are presented to the network. After a random delay, a ‘Set’ signal is flashed, instructing the network to start producing a ramping output. The output time interval *t*_*out*_ is defined as the time elapsed between the ‘Set’ signal and the time point when the output ramp reaches a predetermined threshold value. Bottom: We trained recurrent neural networks (RNNs) where task-relevant inputs (left) are fed to the recurrently connected units. The network output is defined as a linear combination of the activity of recurrent units. **B** Structure of tasks. We considered two tasks: Cue-Set-Go (CSG) and Measure-Wait-Go (MWG). In the CSG task (top), the amplitude of a tonic input cue (grey line), present during the whole trial duration, indicates the time interval to be produced. In the MWG task (bottom), two input pulses are fed to the network (“inp 1” and “inp 2”), followed by a random delay. The input time interval *t*_*in*_ between the two pulses indicates the target output interval *t*_*out*_. **C** Characterizing generalization in terms of input-output functions. We trained networks on a set of training samples (colored squares), so that when learning converges, a specific cue amplitude (top) or input interval *t*_*in*_ (bottom) corresponds to an output interval *t*_*out*_. We then tested how trained networks generalized their outputs to inputs not seen during training, both within (interpolation, grey shaded region) and beyond (extrapolation) the training range, as illustrated with the dotted line. Note the different input axis for the CSG task (top, cue amplitude) and the MWG task (bottom, time interval). **D** To examine the influence of dimensionality on generalization, we compared two types of networks trained on each task. Low-rank networks had a connectivity matrix ***J*** of fixed rank *R*, which directly constrained the dimensionality of the dynamics (Dubreuil et al., 2022). Such networks were generated by parametrizing the connectivity matrix as in Eq. (2), and training only the entries of the connectivity vectors ***m***^(*r*)^ and ***n***^(*r*)^ for *r* = 1 … *R*. Unconstrained networks instead had full-rank connectivity matrices, where all the matrix entries were trained.

We first examined the role of dimensionality for generalization when a tonic input is present. We considered the Cue-Set-Go (CSG) task (Wang et al., 2018), where the goal is to produce an output time interval associated to a tonic input cue, the level of which varies from trial to trial (Fig. 1 B top). Correctly performing this task required learning the association between each cue and target output interval. We found that networks with rank-two connectivity could solve this task with similar performance to unconstrained networks (Fig. 2 A, S1), but led to lower-dimensional dynamics (Fig. 2 A-B top, see Methods). During training, we used four different input amplitudes that were linearly related to their corresponding output intervals (four different color lines in Fig. 2 A). We then examined how the two types of networks generalized to previously unseen amplitudes. We found that both rank-two and unconstrained networks were able to interpolate to cue amplitudes in between those used for training (Fig. 2 C top). However, only low-rank networks showed extrapolation (i.e., a smooth monotonic input-output mapping) when networks were probed with cue amplitudes beyond the training range. Specifically, as the amplitude of the cue input was increased, low-rank networks generated increasing output time intervals (Fig. 2 C, see Fig. S2 for additional examples of similarly trained networks), while unconstrained networks instead stopped producing a ramping signal altogether (see Fig. S3).

**Figure 2:**
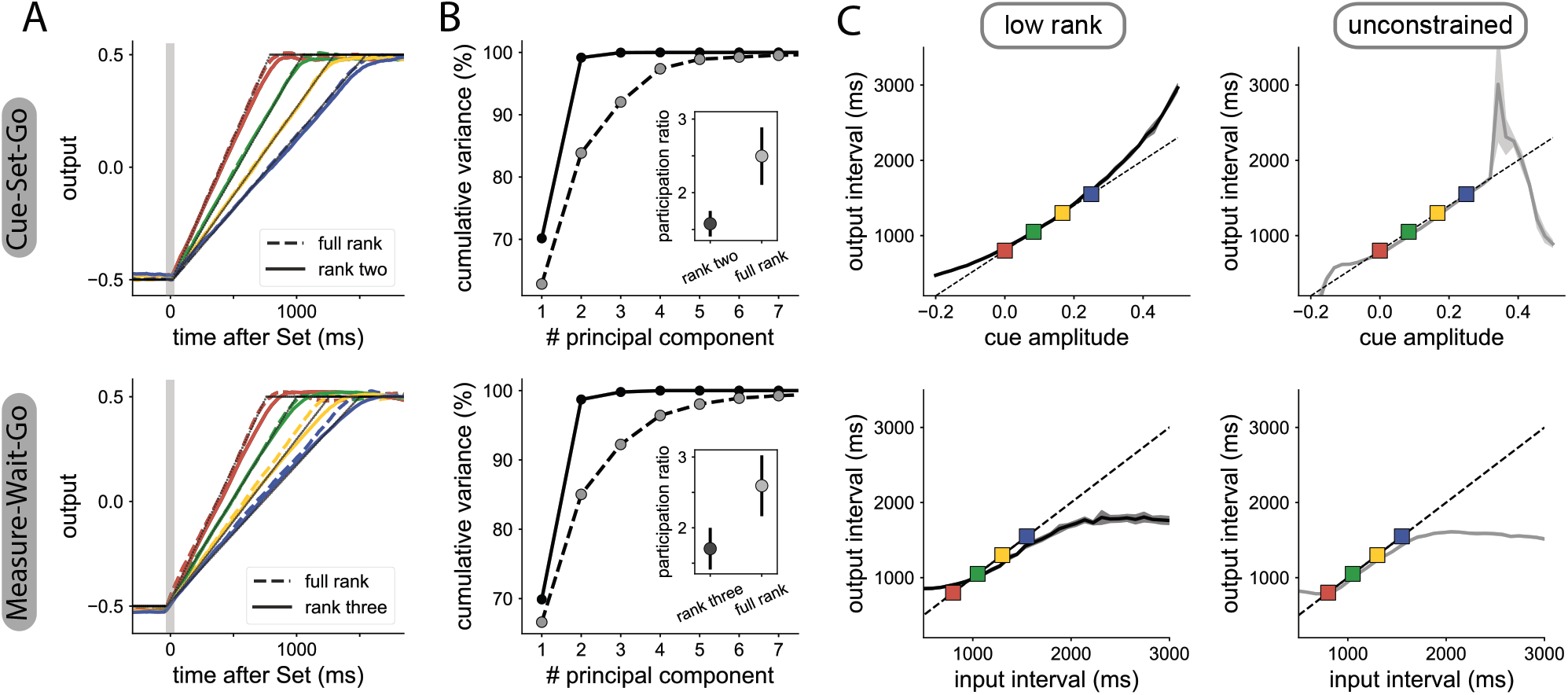
Influence of the dimensionality of network dynamics on generalization. Top: CSG task; bottom: MWG task. **A** Network output of RNNs trained on four different time intervals. RNNs with minimal rank *R* = 2 (solid colored lines) are compared with unconstrained RNNs (dashed colored lines). The thin black lines represent the target ramping output. Note that the dashed colored lines overlap with the solid ones. **B** Dimensionality of neural dynamics following the ‘Set’ signal. Cumulative percentage of explained variance during the measurement epoch as a function of the number of principal components, for rank-two networks (solid line) and unconstrained networks (dashed line). Inset: participation ratio of activity during the whole trial duration (See Methods), with mean and SD over 5 trained RNNs. **C** Generalization of the trained RNNs to novel inputs within and beyond the training range. Low-rank (left) and unconstrained (right) RNNs. Lines and the shaded area respectively indicate the average output interval and the standard deviation estimated over ten trials per cue amplitude. In the CSG task (top row), the rank-two network interpolates and extrapolates, while the unconstrained network does not extrapolate. In the MWG task (bottom row), both types of networks fail to extrapolate to input intervals beyond the training set. The longest output interval that each network can generate is close to the longest output interval learned during training. The delay was randomized during training but the performance plots are based on a fixed delay of 1500ms (see S4 for other delays). See Figs. S2 and S5 for additional trained networks.

We next asked whether low-dimensional dynamics enable generalization in absence of tonic inputs. We therefore turned to a more complex task, Measure-Wait-Go (MWG), in which the desired output was indicated by the time interval between two identical pulse inputs (Fig. 1 B bottom). The network was required to measure that interval, keep it in memory during a delay period of random duration, and then reproduce it after the ‘Set’ signal.

In this task, since the input interval is not encoded as a tonic input, the network must estimate the desired interval by tracking time between the two input pulses. We found that a minimal rank *R* = 3 was required for low-rank networks to match the performance of unconstrained networks (Fig. 2 A-B bottom, Fig. S1). We then examined how low-rank and unconstrained networks responded to input intervals unseen during training. Both types of networks interpolated well to input intervals in between those used for training. However neither the low-rank nor the unconstrained networks were able to extrapolate to intervals longer or shorter than those in the training set (Fig. 2 C bottom, see Fig. S4 for the input-output curves for different delays and Fig. S5 for the same analysis on other trained networks). Instead, the maximal output interval saturated at a value close to the longest interval used for training, thereby leading to a sigmoid input-output function.

Altogether, these results indicate that generalization does not emerge from low-dimensionality alone (MWG task), but depends on the presence of tonic inputs (CSG task).

### 2.2 Combining contextual inputs in low-rank networks enable parametric control of extrapolation

The key difference between the CSG and MWG tasks was that the interval in the CSG was specified by a tonic input, whereas in the MWG task, it had to be estimated from two brief input pulses. Since low-rank networks were able to extrapolate in the CSG but not the MWG task, we hypothesized that the tonic input in CSG may act as a parametric control signal allowing extrapolation in low-rank networks.

To test this hypothesis, we designed an extension of the MWG with a tonic contextual input, which we refer to as the Measure-Wait-Go with Context (MWG+Ctxt). The MWG+Ctxt task was identical to the original MWG task in that the network had to measure an interval from two pulses, remember the interval over a delay, and finally produce the interval after ‘Set’. However, on every trial, the network received an additional tonic input that specified the range of input intervals the network had to process (Fig. 3 A). A low-amplitude tonic input indicated a relatively shorter set of intervals (800-1550 ms), and a higher tonic input, a set of longer intervals (1600-3100 ms). Note that the functional role of this tonic input is different from that of the CSG; in CSG, the input specifies the actual interval, whereas in the MWG+Ctxt task, it only specifies the underlying distribution. The contextual input therefore provides additional information that is not strictly necessary for performing the task, but may facilitate it.

**Figure 3:**
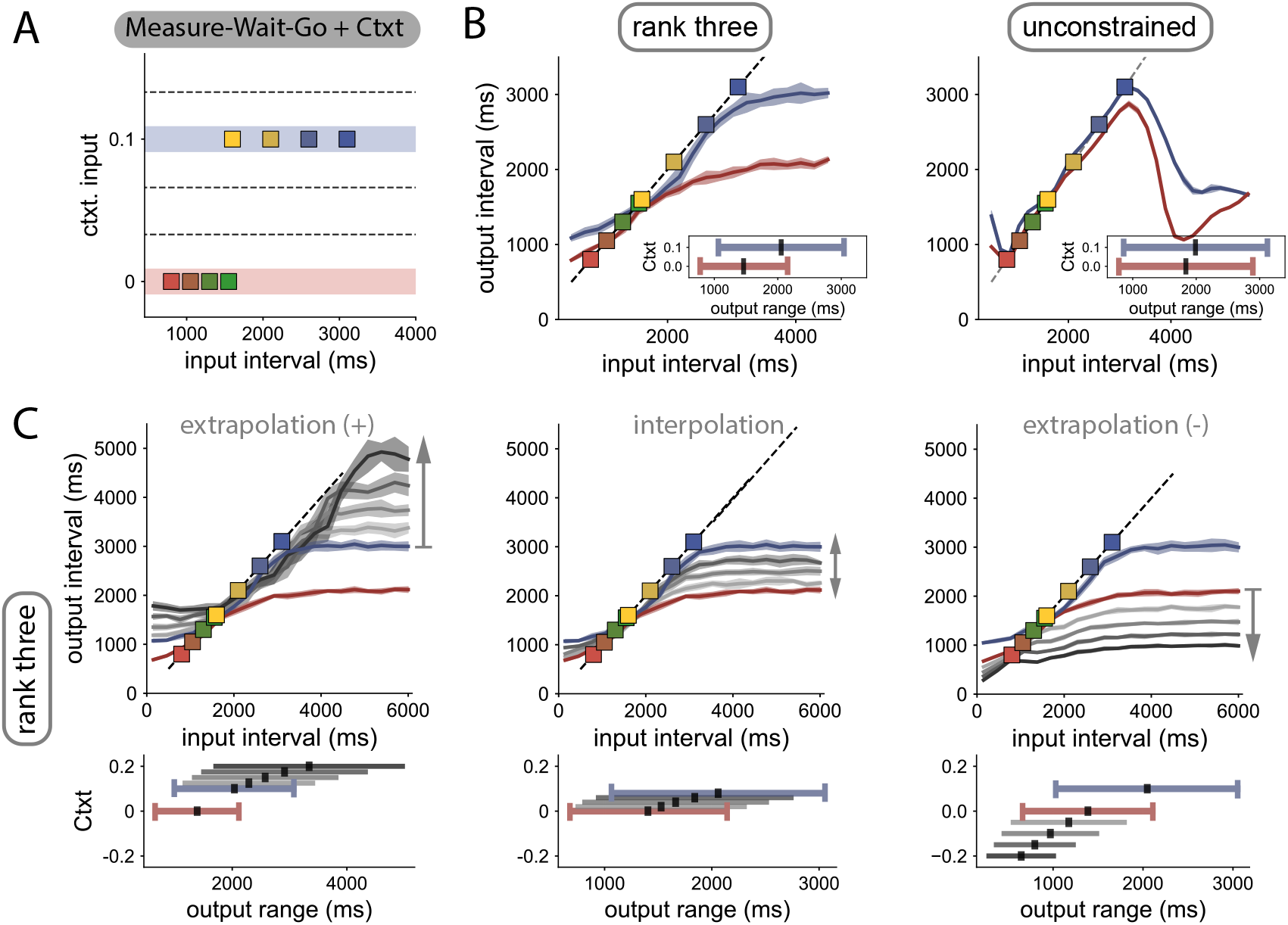
Control and extrapolation in the Measure-Wait-Go task with contextual cues. **A** Task design. Left: Input time intervals *t*_*in*_ used for training (colored squares) are sampled from two distributions. A contextual cue, present during the whole trial duration, indicates from which distribution (red or blue lines) the input interval was drawn. After training, we probed how novel contextual cues (dashed lines) influence the input-output transform of the network. **B** Generalization to time intervals not seen during training, when using the two trained contextual cues (red and blue lines). Contextual cues modulate the input-output function in law-rank networks (left), but not in unconstrained networks (right). Lines and shading indicate average output interval *t*_*out*_ and standard deviation for independent trials. Inset: range of output intervals that the networks can produce in the different contexts. Black lines indicate the midpoint of the range. **C** Generalization in the low-rank network when using contextual cues not seen during training (context values shown in bottom insets). Top: Effects of contextual inputs above (left), within (center) and below (right) the training range. Bottom: Modulation of the range of output intervals as a function of the trained and novel contextual cues (colored and grey lines, respectively). See Fig. S7 for additional low-rank RNNs and Fig. S8 for unconstrained RNNs.

Similar to what we found for the MWG task, both rank-three and unconstrained networks were able to solve the MWG+Ctxt task, and reproduced input intervals sampled from both distributions (Fig. S6). We then examined how each type of network generalized to unseen intervals when associated with one of the two contextual cues used during training (Fig. 3 B).

Building on what we had learned from extrapolation in the CSG task, we reasoned that low-rank networks performing the MWG+Ctxt task may be able to use the tonic input as a control signal enabling generalization to interval ranges outside those the networks were trained for. To test this hypothesis, we examined generalization to both unseen input intervals and unseen contextual cues, comparing as before low-rank and unconstrained networks.

As in the original MWG task, when the input intervals were increased beyond the training range, the output intervals saturated to a maximal value in both types of networks. The only notable difference was that, in low-rank networks, this maximal value depended on the contextual input (Fig. 3 B left), and was set by the longest interval of the corresponding interval distribution, while in unconstrained networks it did not depend strongly on context (Fig. 3 B right). More generally, in low-rank networks, the contextual inputs modulated the input-output function, and biased output intervals towards the mean of the corresponding input distribution. This was evident from the distinct outputs the network generated for identical intervals under the two contextual inputs (Fig. 3 B left), reminiscent of what has been observed in human and monkey behavior in similar tasks (Jazayeri and Shadlen, 2010, 2015; Sohn et al., 2019). In contrast, unconstrained networks were only weakly sensitive to the contextual cue and reproduced all intervals within the joint support of the two distributions similarly (Fig. 3 B right).

We next probed generalization to values of the contextual input that were never presented during training. Strikingly, in low-rank networks we found that novel contextual inputs parametrically controlled the input-output transform, therefore generalizing across context values (Fig. 3 C). In particular, contextual inputs outside of the training range led to strong extrapolation to unseen values of input intervals. Indeed, contextual inputs larger than used for training expanded the range of output intervals beyond the training range and shifted its mid-point to longer values (Fig. 3 C, bottom left), up to a maximal value of the contextual cue that depended on the training instance (Fig. S7). Conversely, contextual cues smaller than used in training instead reduced the output range and shifted its mean to smaller intervals (Fig. 3 C, bottom right). In contrast, in networks with unconstrained rank, varying contextual cues did not have a strong effect on the input-output transform, which remained mostly confined to the range of input intervals used for training (Fig. S8).

Altogether, our results indicate that, when acting on low-dimensional neural dynamics, contextual inputs serve as control signals to parametrically modulate the input-output transform performed by the network, thereby allowing for successful extrapolation beyond the training range. In contrast, unconstrained networks learnt a more rigid input-output mapping that could not be malleably controlled by new contextual inputs.

### 2.3 Non-linear activity manifolds underlie contextual control of extrapolation

To uncover the mechanisms by which low-rank networks implement parametric control for flexible timing, we examined the underlying low-dimensional dynamics (Sussillo and Barak, 2013; Vyas et al., 2020; Dubreuil et al., 2022). In low-rank networks, the connectivity constrains the activity to be embedded in a low-dimensional subspace (Mastrogiuseppe and Ostojic, 2018; Beiran et al., 2021; Dubreuil et al., 2022), allowing for a direct visualisation. Examining the resulting low-dimensional dynamics for networks trained on flexible timing tasks, in this section we show that the neural trajectories are attracted to non-linear manifolds along which they evolve slowly. The dynamics on, and structure of, these manifolds across conditions determine the extrapolation properties of the trained networks. Here we summarize this reverse-engineering approach and the main results. Additional details are provided in the Methods section.

#### Low-dimensional embedding of neural activity

The RNNs used to implement flexible timing tasks consisted of *N* units, with the dynamics for the total input current *x*_*i*_ to the *i*-th unit given by:

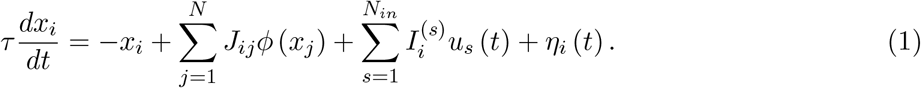

Here *ϕ* (*x*) = tanh (*x*) is the single neuron transfer function, *J*_*ij*_ is the strength of the recurrent connection from unit *j* to unit *i*, and *η*_*i*_ (*t*) is a single-unit noise source. The networks received *N*_*in*_ task-dependent scalar inputs *u*_*s*_ (*t*) for *s* = 1, …, *N*_*in*_, each along a set of feed-forward weights that defined an *input vector* 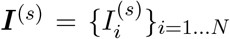 across units. Inputs *u*_*s*_ (*t*) were either delivered as brief pulses (‘Set’ signal, input interval pulses in the MWG task) or as tonic signals that were constant over the duration of a trial (cue input in the CSG task, contextual input in the MWG+Ctxt task).

The collective activity in the network can be described in terms of trajectories 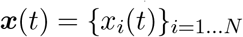 in the *N*-dimensional state space, where each axis corresponds to the total current received by one unit (Fig. 4 A). In low-rank networks, the connectivity constrains the trajectories to reside in a lowdimensional linear subspace of the state space (Mastrogiuseppe and Ostojic, 2018; Beiran et al., 2021; Dubreuil et al., 2022), that we refer to as the *embedding space* (Jazayeri and Ostojic, 2021). In a rank-*R* network, the recurrent connectivity can be expressed in terms of *R* pairs of *connectivity vectors* 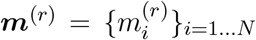 and 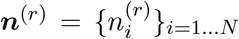 for *r* = 1 … *R*, which together specify the recurrent connections through

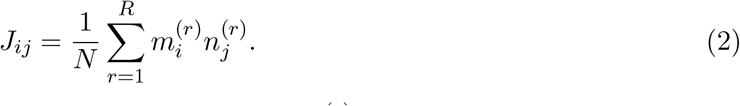

**Figure 4:**
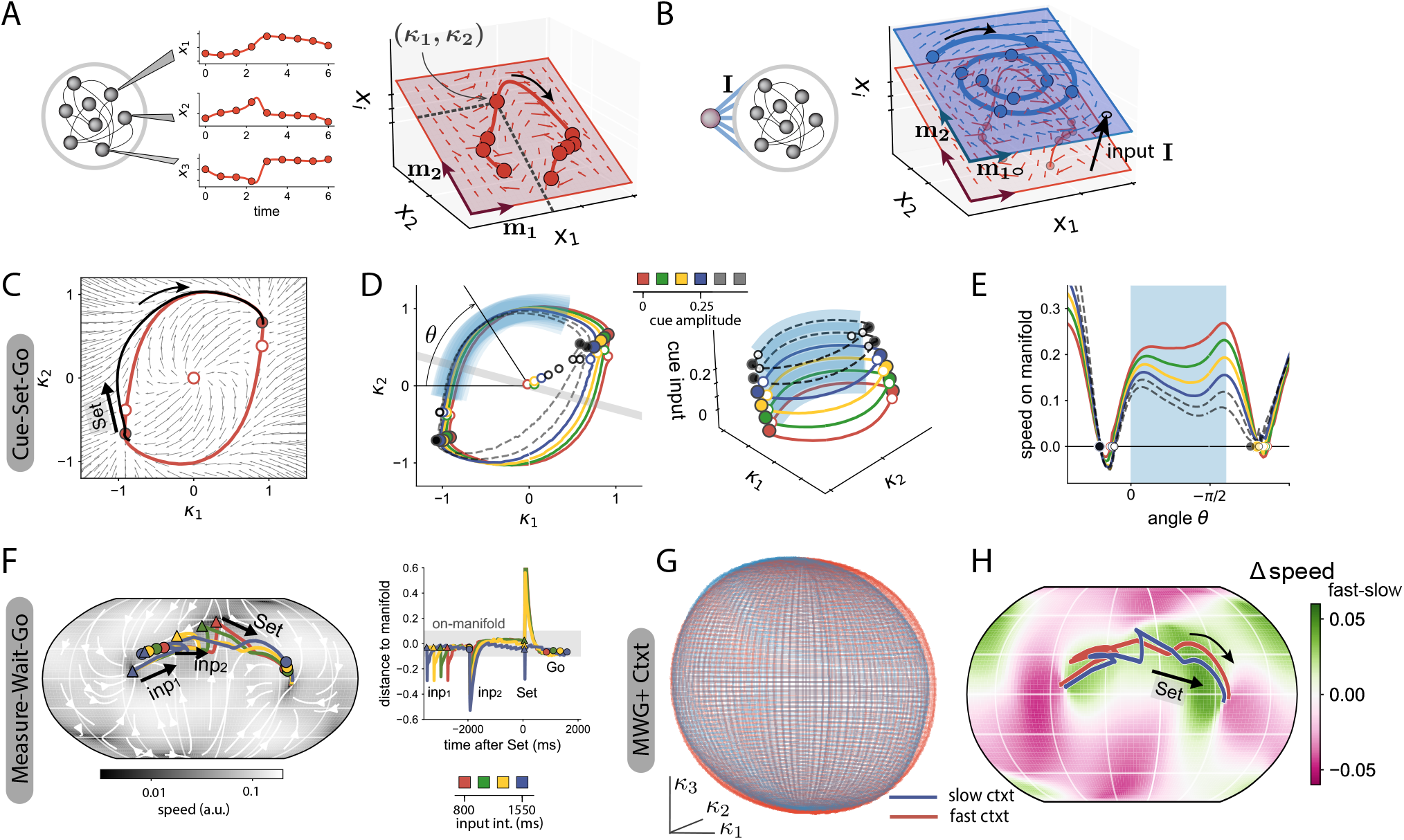
Reverse-engineering RNNs trained on flexible timing tasks. **A-B** Low-dimensional description of activity dynamics. **A** The temporal activity of different units in the network (left) can be represented as a trajectory in state space (right, thick red line), where each axis corresponds to the activity *x*_*i*_ of one unit. In low-rank networks, trajectories are embedded in a lower dimensional space (red shaded plane, see Eq. (3)). For instance, in a rank-two network, in absence of input, this embedding space is spanned by connectivity vectors ***m***^(1)^ and ***m***^(2)^, so that neural trajectories can be parametrized by two recurrent variables *κ*_1_ and *κ*_2_, and their dynamics represented in terms of a flow field on the embedding space (arrows, see Eq. (4)). **B** External inputs increase the dimensionality of the embedding space. In this illustration, a single input is added, so that the embedding subspace is now three-dimensional: two dimensions for the recurrent subspace, spanned by ***m***^(1)^ and ***m***^(2)^, and one additional dimension for the input vector ***I***. If the input is tonic, it shifts the recurrent subspace (***m***^(1)^ - ***m***^(2)^) along the input direction (black arrow, from the red to the blue plane) and modifies the flow fields. **C-E** RNN trained on the CSG task. Color-filled dots represent stable fixed points, and white dots represent unstable fixed points. **C** Dynamical landscape in the embedding subspace of trained rank two network (small arrows), and neural trajectory for one trial (black line, *u*_*cue*_ = 0). The red line represents the non-linear manifold to which dynamics converge from arbitrary initial conditions (Fig. S9). The trial starts at the bottom left stable fixed point. The ‘Set’ pulse initiates a trajectory towards the opposite stable fixed point, which quickly converges to the slow manifold (see Fig. S11 A), and evolves along it. **D** Manifolds generated by a trained RNN for different amplitudes of the cue. Colored lines correspond to the different cues used for training and black dashed lines correspond to amplitudes beyond the training range. Left: Two-dimensional projections of the manifolds onto the recurrent subspace spanned by vectors ***m***^(1)^ and ***m***^(2)^. The grey line is the projection of the readout vector on the recurrent subspace. Right: Three dimensional visualization of the manifolds in the full embedding subspace, spanned by ***m***^(1)^, ***m***^(2)^ and ***I***_***cue***_. The shaded blue region indicates the section of the manifolds along which neural trajectories evolve when performing the task. Any state on the manifold can be determined by the polar coordinate *θ*. Increasing the cue amplitude, even to values beyond the training range (black dashed lines), keeps the geometry of manifolds invariant. **E** Speed along the manifold as a function of the polar angle *θ*, that parametrizes all on-manifold states. The speed along the manifold is scaled by the cue amplitude, even beyond the training range. Shaded blue region as in **D. F** RNN trained on the MWG task. Left: The dynamics in the embedding subspace of the trained rank-three network generated a spherical manifold to which trajectories were quickly attracted after fast transient responses to input pulses (Fig. S9). Right: distance of trajectories solving the task to the manifold’s surface. **G** RNN trained on the MWG+Ctxt task. Spherical manifolds on the recurrent subspace for two different contextual cues (red: fast context, *u*_*ctxt*_ = 0, blue: slow context, *u*_*ctxt*_ = 0.1). **H** The contextual cue modulates the speed of dynamics along the manifold. The colormap shows the difference in speed on the manifold’s surface between the two contexts. In the region of the manifold where trajectories evolve (red and blue lines), the speed is lower for stronger contextual cues (slow context). A stronger contextual amplitude slows down the non-linear manifold in the region of interest (see Fig. S11 C for novel contextual amplitudes).

The connectivity vectors, as well as the feed-forward input vectors ***I***^(*s*)^, define specific directions within the *N*-dimensional state space that determine the network dynamics. In particular, the trajectories are confined to the embedding space spanned by the recurrent connectivity vectors ***m***^(*r*)^ and the feed-forward input vectors ***I***^(*s*)^ (Fig. 4 A). The trajectories of activity ***x***(*t*) can therefore be parameterized in terms of Cartesian coordinates along the basis formed by the vectors ***m***^(*r*)^ and ***I***^(*s*)^ (Mastrogiuseppe and Ostojic, 2018; Beiran et al., 2021; Dubreuil et al., 2022):

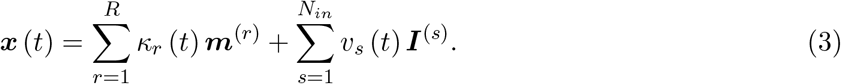

The variables 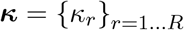 and 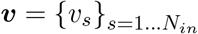 represent respectively activity along the *recurrent* and input-driven subspaces of the embedding space (Wang et al., 2018). The dimensionality of the embedding space is therefore *R* + *N*_*in*_, where *R* dimensions correspond to the recurrent subspace, and *N*_*in*_ to the input-driven subspace. The evolution of the activity in the network can then be described in terms of a dynamical system for the recurrent variables ***κ*** driven by inputs ***v*** (Beiran et al., 2021; Dubreuil et al., 2022):

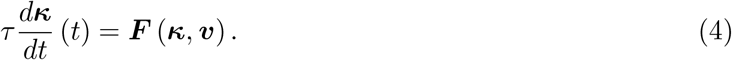

In Eq. (4), for constant inputs the non-linear function ***F*** describes the dynamical landscape that determines the *flow* of the low-dimensional activity in the recurrent subspace (Fig. 4 A). Pulse-like inputs instantaneously shift the position of activity in the embedding space, such that the transient dynamics in between pulses are determined by this dynamical landscape. Instead, tonic cue inputs, such as the signals used in the CSG and MWG+Ctxt tasks, shift the recurrent subspace within the embedding space and thereby modify the full dynamical landscape (Fig. 4 B). The low-dimensional embedding of neural trajectories can therefore be leveraged to explore the dynamics of trained recurrent neural networks, by directly visualizing the dynamical landscape and the flow of trajectories in the recurrent subspace instead of considering the full, high-dimensional state space.

#### Non-linear manifolds within the embedding space

In low-rank networks, the trajectories of activity reside in the low-dimensional embedding space, but they do not necessarily explore that space uniformly. Examining networks trained on flexible timing tasks, we found that neural trajectories quickly converged to lower-dimensional, non-linear regions within the embedding space. We refer to these regions as *non-linear manifolds* (Jazayeri and Ostojic, 2021), and we devised methods to identify them from network dynamics (Methods and Fig. S9). We next describe these manifolds and their influence on dynamics and computations for networks trained on individual tasks.

#### Cue-Set-Go task

For rank-two networks that implemented the CSG task, the recurrent subspace was of dimension two and was parameterized by Cartesian coordinates *κ*_1_ and *κ*_2_ (Fig. 4 C). In a given trial, corresponding to a fixed cue amplitude, we found that the dynamical landscape on this recurrent subspace displayed an attractive ring-like manifold on which the dynamics were slow (Fig. 4 C, red line). This manifold connected two stable fixed points through two intermediate saddle points (Fig. 4 C, red and white dots respectively). At trial onset, the neural activity started at one first stable fixed point, and was then pushed by the ‘Set’ pulse above a saddle point. This generated a neural trajectory in the recurrent subspace (Fig. 4 C, black line) that was quickly attracted to the ring-like manifold (Figs. S9 and S11), and subsequently followed slow dynamics along this manifold towards the second stable fixed point. The position on this non-linear manifold therefore represented the time since the ‘Set’ pulse, and was directly transformed into a ramping output by the readout of the network.

Different trials in the CSG task correspond to different amplitudes of the tonic input cue that shift the position of the recurrent subspace within the embedding space and modulate the dynamical landscape on it (Fig. 4 D). For each value of the cue amplitude, we found that trajectories evolved along parallel ring-like manifolds, which together formed a two-dimensional cylinder when visualised in the three-dimensional embedding space defined by *κ*_1_, *κ*_2_ and the input cue as a third coordinate (Fig. 4 D right). The shape of the ring-like manifolds and the position of fixed points were largely invariant, but the amplitude of the cue controlled the speed of the dynamics along each ring (Fig. 4 E), and thereby the slope of the ramping output that determined the output interval (Fig. 2 A). Because of the cylindrical geometry, extending cue amplitudes to values outside of the training range preserved the overall structure of the manifold (Fig. 4 D blacked dashed line), and thereby ensured lawful extrapolation of the required outputs. The modulation of speed along a low-dimensional manifold of invariant geometry therefore subserves the contextual control of extrapolation in the CSG task.

#### Measure-Wait-Go task

For rank three networks that implemented the MWG task, the recurrent subspace was three-dimensional. Within that subspace, we found that the trajectories were attracted to a spherically-shaped manifold, only diverging from its surface during the fast transient responses to the input pulses (Fig. 4 F, Fig. S9).

The dynamics of activity underlying the basic MWG task could therefore be described in terms of trajectories along a non-linear manifold, in a manner analogous to individual trials in the CSG task. During the measurement period, the two input pulses that defined the input time interval led to trajectories that quickly converged to a localized region on the spherical manifold where the speed was minimal (Fig. S11). This region played the role of a line attractor, the position along which encoded the input time interval during the delay period. The subsequent ‘Set’ input then generated trajectories that evolved towards the other side of the sphere, at speeds set by the initial condition on the line attractor, therefore leading to ramp signals with varying slopes (Fig. S3). Crucially, the line attractor that encoded the input interval occupied a bounded region on the sphere, so that any input interval outside of the training range converged to one of the extremities of the attractor. Thus, the finite limits of the line attractor lead to a sigmoidal input-output function, where the output intervals were saturated to the bounds of the training range as seen in Fig. 2 C.

For the extended MWG task with a contextual cue, the additional tonic contextual inputs modified the flow of the dynamics in the three-dimensional recurrent space. In a manner analogous to the CSG task, increasing the amplitude of the contextual input largely preserved the shape and position of the attractive spherical manifold (Fig. 4 G), while scaling the speed of the dynamics on it (Fig. 4 H). The contextual input thereby controlled the speed of trajectories of activity and parametrically modulated the input-output transform performed by the network. This effect extended to values of contextual inputs well beyond the trained region (Fig. S9), and thereby controlled extrapolation to previously unseen values for both input intervals and contextual inputs.

In summary, reverse-engineering low-rank networks trained on the CSG and MWG tasks revealed that in both tasks extrapolation beyond the training range relied on a mechanism based on an invariant geometry of underlying neural activity manifolds, the dynamics along which were parametrically controlled by tonic inputs.

### 2.4 Controlling the geometry and dynamics on non-linear manifolds

Our reverse-engineering analysis revealed that parametric control of extrapolation in trained low-rank networks relied on modulating dynamics over non-linear manifolds of invariant geometry. To further unravel how the properties of recurrent connectivity and tonic inputs respectively contribute to controlling the geometry of, and dynamics on non-linear activity manifolds, we next investigated simplified, mathematically tractable networks. In such networks, a mathematical analysis allows us to directly infer dynamics from the connectivity and input parameters and to synthesize networks that perform specific computations (Mastrogiuseppe and Ostojic, 2018; Schuessler et al., 2020a; Beiran et al., 2021; Dubreuil et al., 2022; Pollock and Jazayeri, 2020), thereby demonstrating that the mechanisms identified through reverse-engineering are sufficient to implement generalization in flexible timing tasks.

We considered the restricted class of Gaussian low-rank RNNs (Mastrogiuseppe and Ostojic, 2018; Beiran et al., 2021; Dubreuil et al., 2022) where for each neuron *i*, the set of components 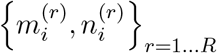 and 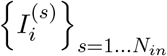 along connectivity and input vectors are drawn randomly and independently from a zero-mean multivariate Gaussian distribution (Fig. S10 A). Unlike trained low-rank networks, which in principle depend on a high number of parameters (order *N*) these simplified networks are fully specified by a few parameters (order (2*R* + *N*_*in*_)^2^) that correspond to the entries of the covariance matrix of the underlying Gaussian distribution, or equivalently the matrix of pairwise *overlaps* between connectivity and input vectors (Fig. S10 A right, Methods). We therefore investigated how this overlap structure determines the dynamics that underlie computations.

#### Generating slow manifolds through recurrent connectivity

We first focused solely on recurrent interactions, and investigated which type of overlap structure between connectivity vectors generates slow attractive manifolds, as seen in trained networks. In particular, we analyzed how the position of the fixed points and speed along manifolds depend on the overlap matrix. As reported in previous studies (Mastrogiuseppe and Ostojic, 2018; Pereira and Brunel, 2018; Beiran et al., 2021) and detailed in the Methods, a mean-field analysis shows that in the limit of large networks, attractive manifolds arise when the overlap matrix is diagonal and its non-zero entries are equal to each other and sufficiently strong (Fig. S10 B left). In Gaussian low-rank RNNs, such an overlap matrix, proportional to the identity, implies that the dynamics induced by recurrent connectivity are symmetric along the different directions of the recurrent subspace spanned by the connectivity vectors ***m***^(*r*)^ for *r* = 1 … *R*. In general, for a rank R network, such a symmetry induces an attractive *R*-dimensional spherical manifold in neural space where each point on the manifold is an attractive fixed point of the dynamics (see Methods). Accordingly, for rank-two networks this structure leads to a continuous ring attractor, embedded in a plane (Fig. S10 B), and for rank-three networks to a spherical attractor.

The mean-field analysis formally holds in the limit of infinitely large networks. Dynamics in networks of finite size can be described by adding random perturbations to the overlap matrix describing the connectivity, which is therefore never perfectly diagonal. Perturbations away from a diagonal overlap structure break up the continuous attractor so that only a small number of points on it remain exact fixed points of the dynamics (Fig. S10 C, see Methods for details). On the remainder of the original continuous attractor, the flow of the dynamics however stays very slow even for relatively large deviations from a diagonal overlap structure. For rank two connectivity, the continuous attractor predicted by the mean-field analysis therefore leads to a ring-like manifold on which slow dynamics connect stable fixed points and saddles (Fig. S10 C, middle), as seen in trained low-rank networks (Fig. 4 C-D). Specific deviations from a diagonal overlap matrix moreover determine the precise position of the fixed points on the manifold. In particular, a weak off-diagonal component, corresponding to a non-zero overlap between connectivity vectors of different rank-one structures, rotates the position of saddle points and brings them closer to stable fixed points (Fig. S10 D). This type of structure leads to long transient trajectories from a saddle to a fixed point, analogous to those underlying ramping signals in trained networks (Fig. 4 C).

#### Controlling dynamics through tonic inputs

We next examined how a tonic cue, i.e. a constant external input along an input vector ***I***, impacts the geometry of recurrently-generated manifolds and the dynamics on them. In particular, we studied what properties of the input vectors and recurrent connectivity produce a change in dynamics such as the one observed in the CSG task (Fig. 4 D) landscape. Our mean-field analysis showed that the geometrical arrangement between the input vector ***I*** and recurrent vectors, as quantified by their overlaps, was the key factor that determined the effect of the input on the manifold. We therefore distinguished between *non-specific inputs* for which the input vector was orthogonal to all connectivity vectors, and *subspace-specific inputs*, for which the input vector was correlated with the recurrent connectivity structure.

Non-specific inputs modify both the dynamics on the manifold, and its geometry in neural state space (Fig. 5 A-B, top row). In particular, increasing the amplitude of non-specific inputs shrinks the radius of the activity manifold, until it eventually collapses. In contrast, for subspace-specific inputs, varying the input amplitude modulates the speed of the dynamics (similar to non-specific inputs, Fig. 5 B), but importantly, keeps approximately invariant the geometry of the manifold and the position of fixed points on it (Fig. 5 A bottom). Subspace-specific inputs therefore reproduce the mechanism of speed modulation on invariant manifolds found when reverse-engineering trained networks. Altogether, we found that, although both specific and non-specific tonic inputs can modulate the speed along similar ranges (Fig. 5 B), only specific inputs keep invariant the geometry of the non-linear manifold (Fig. 5 A).

**Figure 5:**
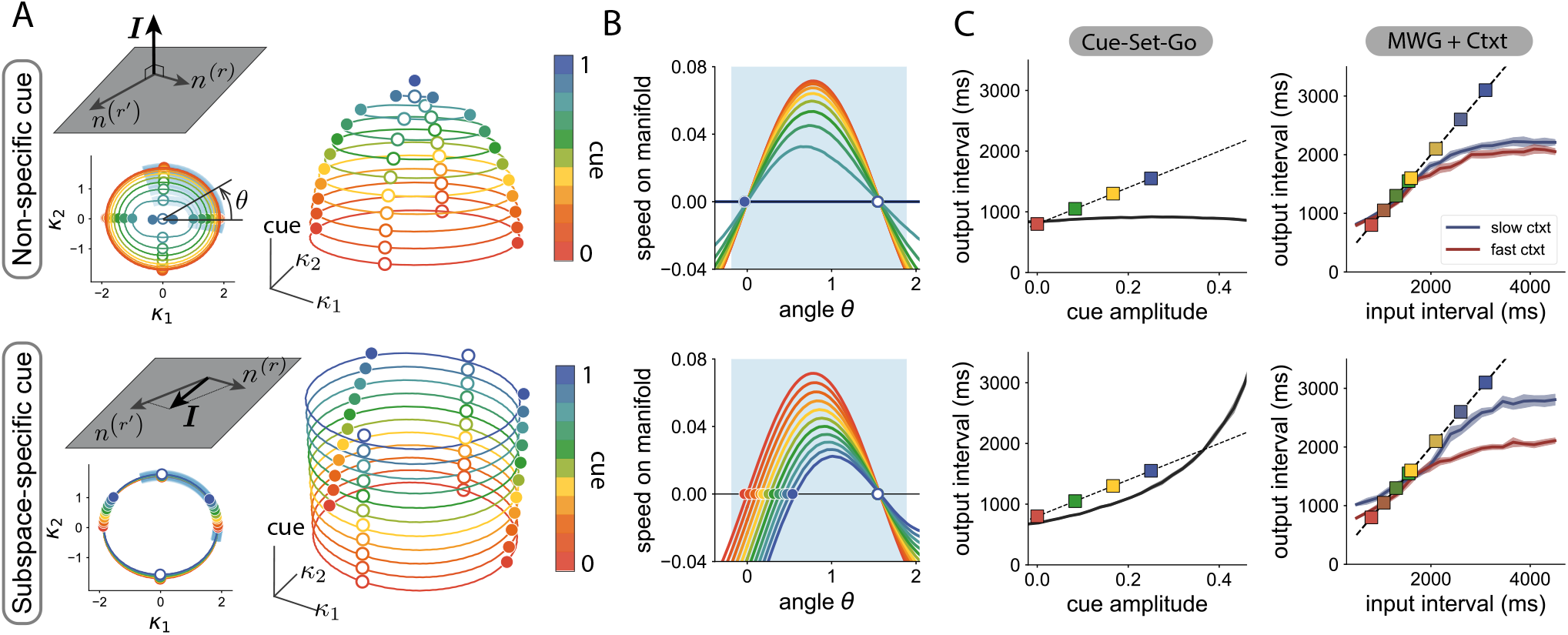
Effect of tonic cue inputs on the geometry of, and dynamics on, manifolds. Top: non-specific cue orthogonal to the recurrent connectivity vectors. Bottom: specific cue input along the subspace spanned by recurrent connectivity vectors. **A** Projection of manifolds on the recurrent subspace (bottom left inset), and on the three-dimensional embedding subspace (right). Different colors correspond to different cue amplitudes. Solid lines indicate the manifold location. Colored dots are stable fixed points, white dots are saddle points. **B** Speed along the manifold, parametrized by the polar angle, along the shaded blue section indicated in **A. C** Performance in the CSG (left) and MWG+Ctxt tasks (right), when either only non-specific (top) or only subspace-specific (bottom) components of the cue of the trained RNN are kept after training. Parameters: A, 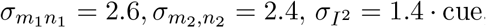, B: 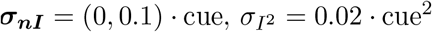.

Based on the mechanism identified using the mean-field analysis, we hypothesized that the input components required to produce flexible timing behavior are those specific to the recurrent subspace. We tested this prediction on the low-rank neural networks trained on the Cue-Set-Go task and Measure-Wait-Go task with context. We perturbed the trained input vectors in two ways; we either kept only input components specific to the connectivity subspace and removed all others, or kept only input components orthogonal to the connectivity subspace. As predicted, keeping only the non-specific components of the input vectors completely hindered the computation (Fig. 5 C top). In contrast, removing all non-specific components from the input vectors in the trained networks did not strongly affect the performance of the network (Fig. 5 C bottom). These results show that subspace-specific input vectors were required to solve timing tasks.

The mean-field analysis of simplified, Gaussian low-rank networks, allowed us to synthesize networks in which tonic inputs control dynamics over non-linear manifolds of invariant geometry. Going one step further, we next capitalized on these insights to directly design networks that perform the Cue-Set-Go and Measure-Wait-Go tasks based on this mechanism, by setting connectivity parameters without training (Fig. S12). Such minimal network models demonstrate that the mechanisms identified by reverse-engineering trained networks are indeed sufficient for implementing the flexible timing tasks over a large range of inputs.

### 2.5 Signatures of the computational mechanism in neural data

Our analyses of network models point to a putative mechanism for rapid generalization in timing tasks. Specifically, in our networks, generalization emerges when inputs are paired with a low-dimensional contextual signal which is parametrically adjusted to the range of possible inputs. To test for the presence of this computational strategy in the brain, we turned to empirical data recorded during flexible timing behavior. We analyzed neural population activity in the dorsomedial frontal cortex (DMFC) of monkeys performing the ‘Ready-Set-Go’ (RSG) task, a time interval reproduction task analogous to the MWG+Ctxt task but without a delay period (Sohn et al., 2019; Meirhaeghe et al., 2021). In this task, animals had to measure and immediately reproduce different time intervals. The feature that made this task comparable to the MWG+Ctxt task was that the time interval on each trial was sampled from one of two distributions (i.e., a “fast” and a “slow” context), and the color of the fixation spot throughout the trial explicitly cued the relevant distribution, thereby providing a tonic contextual cue.

Previous studies have reported on several features of neural dynamics in this task (Sohn et al., 2019; Meirhaeghe et al., 2021). In particular, it was found that during the measure of the intervals, population activity associated with each context evolves along two separate trajectories running at different speeds. While this observation is qualitatively consistent with the contextual speed modulations seen in our network models, it is by itself insufficient to validate the hypothesized mechanism, a tonic contextual input controlling the low-dimensional dynamics. To firmly establish the connection between our model and the neural data, we thus devised a new set of quantitative analyses aimed at directly comparing their dynamics (Remington et al., 2018a).

To guide our comparison between the model and the data, we focused on three main signatures of the low-rank networks. Our first observation was that the neural trajectories associated with the two contexts remain separated along a relatively constant dimension throughout the measurement epoch of the task. Indeed, as we have shown above, the contextual input translates the manifold of activity along a “context dimension” that is orthogonal to the subspace in which the trajectories evolve (Fig. 6 A top left). This translation was also visually manifest in the data: the empirical trajectories associated with the two contexts appeared to evolve in parallel, as shown by a 3D projection of the dynamics (Fig. 6 A bottom left). To rigorously assess the degree of parallelism of the trajectories in higher dimensions, we computed the angle between the vectors separating the two trajectories at different time points (Methods). We expected this angle to be small if the dimension separating the trajectories remained stable over time. Consistent with our initial observations, we indeed found that in the low-rank networks this angle remained close to zero (Fig. 6 A top right). When we performed the same analysis on the neural data, we found a similar degree of parallelism between the trajectories (Fig. 6 A bottom right): in both the model and the data, the angle remained far from 90 degrees for at least 500 ms (Fig. S13 A), indicating that the context dimension was largely stable over time.

**Figure 6:**
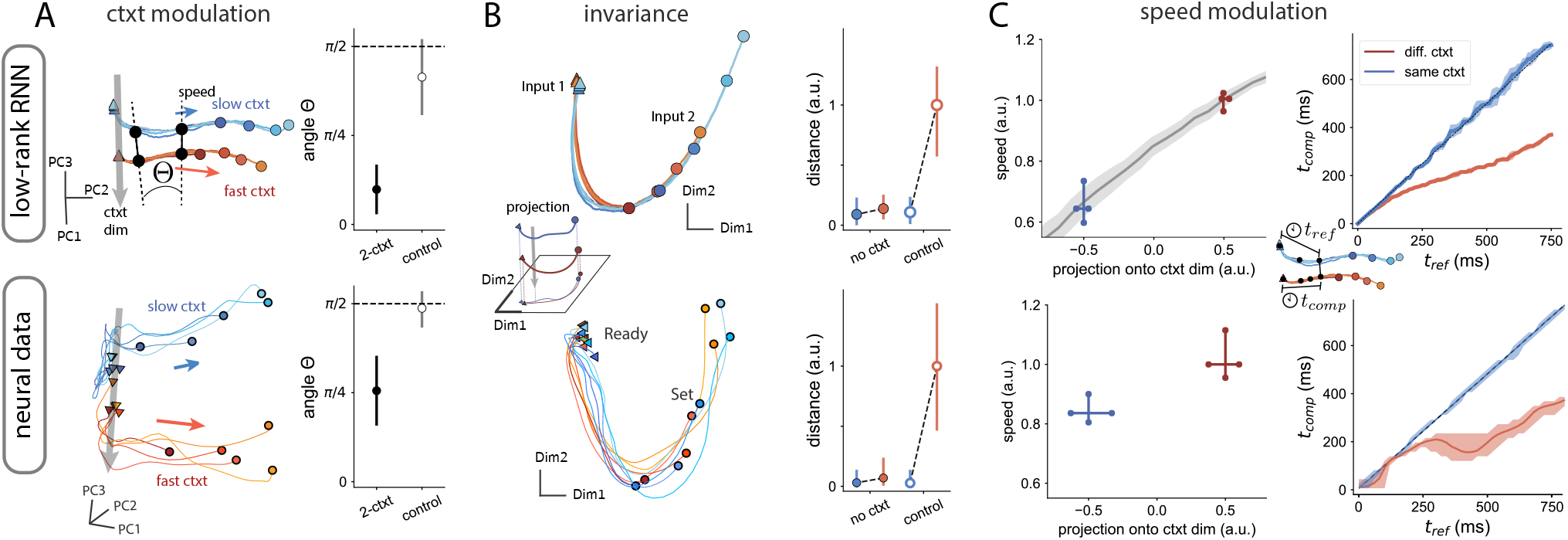
Signatures of the computational mechanism in neural data. Comparison between a low-rank network trained on the MWG+Ctxt task (top) and neural data in a time-reproduction task (Sohn et al., 2019; Meirhaeghe et al., 2021) (bottom). **A** Left. Projections on the first principal components of the activity during the measurement epoch between the two input pulses. Different trajectories correspond to different input intervals; colors denote the two contexts. Right. Mean (in absolute value) and standard deviation of the angle between vectors separating trajectories. Control corresponds to the same analysis with shuffled context categories. **B** Left. Two-dimensional projection of activity orthogonal to the context axis. Trajectories evolve along a non-linear manifold that is invariant across contexts. Right. Average distance between trajectories during the measurement epoch after removing the projection onto context dimension. The distance is computed between each average context trajectory (2-ctxt, red: fast ctxt, blue: slow ctxt) and the reference trajectory, which we chose to be the slow context. The estimate of the distance for the reference trajectory (blue) with itself uses different subsets of trials, as a way to quantify the noise. The control consists of removing the projection of the activity on a random direction (see Methods). **C** Left. Average speed of trajectories during the measurement epoch, as a function of the projection onto the context dimension. In the RNN, the grey line shows the same analysis for context amplitudes not seen during training. There is a monotonic relation between the speed of neural activity during measurement and the projection of the activity onto the context dimension. Right. Speed estimation shown as the time elapsed until reaching a reference state (given by the average trajectory in the slow context). Trajectories from the context different from reference evolve faster after an initial transient at equal speeds. Errorbars indicate 99-confidence interval.

Our second observation was that the neural trajectories in the low-rank networks were invariant across contexts and overlapped in the recurrent subspace (Fig. 6 B left). We visually found a similar invariance in the neural data, after projecting out the context dimension (Fig. 6 B bottom left). To quantify this effect, we calculated the distance between trajectories once the projection of the activity along the previously identified context dimension was removed (see Methods). For comparison, we also calculated the distance between trajectories when removing the projection on a random dimension of the state space (see Fig. S13 B for the time-resolved distances). We found that, both in the networks and the data, the distance was largely reduced to zero once the context dimension was removed (Fig. 6 B right), thus demonstrating a similar degree of invariance in the dynamics.

A final key signature of the low-rank networks was that the speed of dynamics during measurement depended on context: faster in the fast context, slower in the slow context. We measured speed by calculating the amount of change in neural activity in small periods of time. In parallel, we projected the activity onto the context dimension estimated at the beginning of measurement. We found that there was a monotonic relation between the activity along the context dimension and the speed of trajectories, based on the slow and fast contexts (blue and red crosses, Fig. 6 C right). In addition, in the low-rank networks, we were able to establish the full functional relation between speed and projection on context for context amplitudes not seen during training (grey line, Fig. 6 C right).

Direct neural speed estimates are however a measure sensitive to the noise level of the activity. To better compare neural speed in the network and the recordings, we used “kinematic analysis of neural trajectories” (KiNeT), which provides a finer-grain analysis based on relative time elapsed to reach a certain state (see Methods, Remington et al. (2018a)). Using this technique first on the networks, we confirmed that there are strong speed modulations between different contexts. Furthermore, speed modulations only appeared once the transient signal from the beginning of measurement fades out, roughly 200 ms following Ready (Fig. 6 C top). Applying the same technique to the neural data, we observed that the empirical trajectories also diverged in terms of speed only after about 200 ms following the Ready signal (Fig. 6 C bottom).

Altogether, these quantitative analyses show that neural activity exhibited the key signatures of parametric control by a low-dimensional input predicted from computational modeling and theory, suggesting that the dynamics of neural trajectories in the cortex may be optimized to facilitate generalization.

### 2.6 Adaptation to changing input statistics

Our comparison of the network model with neural data argues that flexible timing across contexts is controlled by a tonic input that flexibly modulates the network’s dynamics. So far, we focused on the situation where the two contexts were explicitly cued and well-known to the animals, and we assumed that the corresponding level of the control input was instantaneously set by an unspecified mechanism. A key question is however how the value of this control input can be adaptively adjusted following uncued changes in input data. In this section, we propose a simple mechanism to dynamically update the contextual input based on the input interval statistics, and compare the predictions of this augmented model with behavioral and neural data from a new monkey experiment.

To allow the model to infer changes in context from the recent history of measured intervals, we hypothesize that the network continuously represents and updates an estimate of the mean of the interval distribution. Indeed, previous work has shown that adjusting neural dynamics based on the mean interval may provide the substrate for adapting to new interval statistics (Meirhaeghe et al., 2021). Inspired by this result, we assume that the amplitude of the contextual input linearly encodes the mean input interval, and is adjusted on every trial based on a “prediction error”, i.e. the difference between the measured interval and the current estimate of the mean (Fig. 7 A). The adjustment of the contextual input can thus be formulated as a simple auto-regressive process that continuously tracks the statistics of the intervals (see Methods).

**Figure 7:**
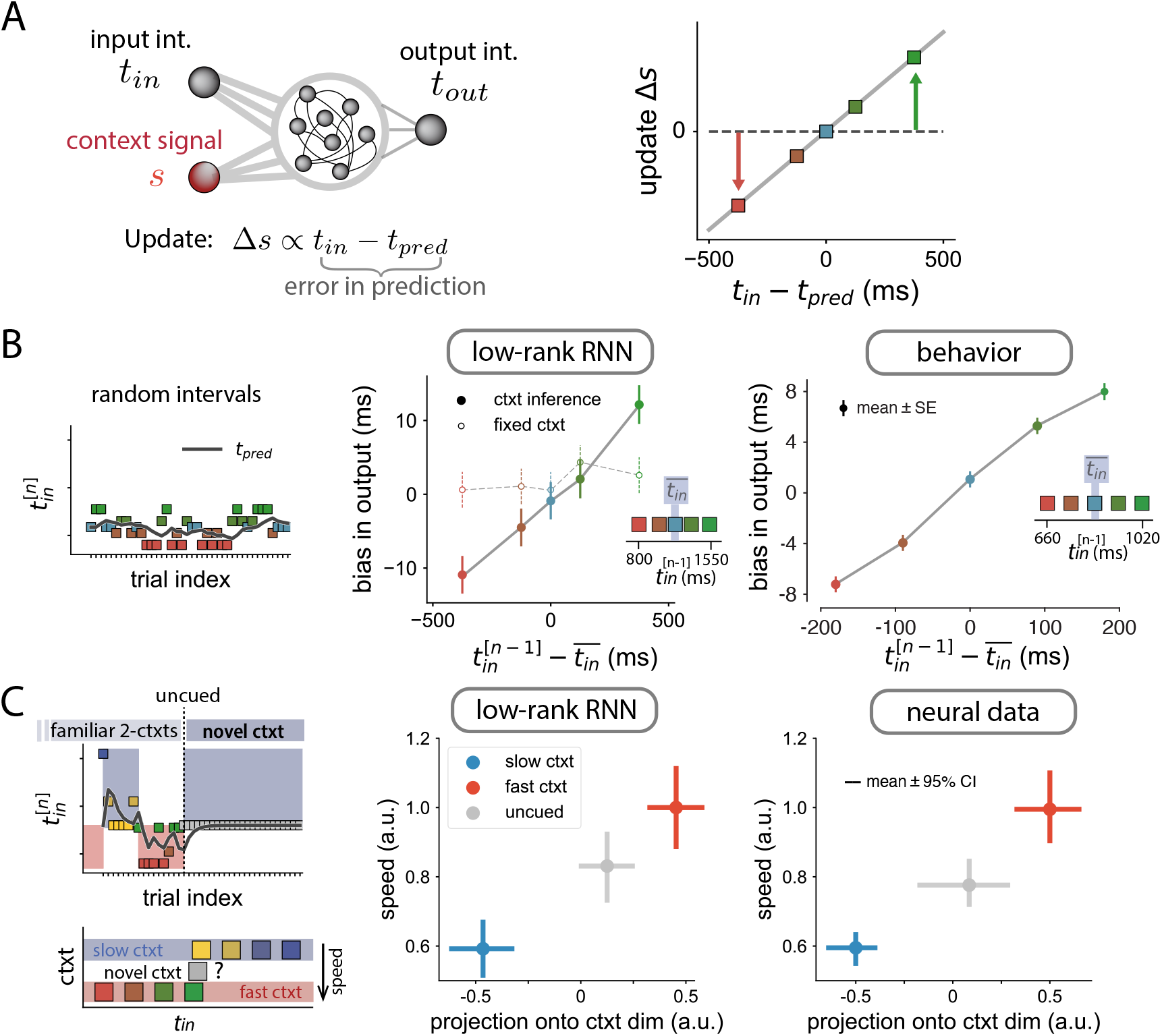
Adaptation to a change in statistics of input intervals. **A** Extended model, in which the context signal amplitude is updated in each trial proportionally to the difference between the input interval and the predicted input interval *t*_*pred*_ (see Methods). **B** Effect of contextual input adjustments in a single-context task. Left. Task structure of a session: randomly interleaved trials from a single distribution. Black line indicates the predicted input interval at every trial. Center. Bias in the output interval of a trained RNN as a function of the interval presented in the previous trial (colored dots). The control data correspond to the same trained RNN with constant context signal. Straight lines connecting the dots were added for visualization. Right. Behavioral bias in monkeys performing a timing task. Insets show the distribution of input intervals, highlighting its mean value, 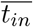. **C** Effect of contextual input adjustments in a two-context adaptation task.Left. Task structure of a session: trials are structured into alternating blocks. In each block, trials are randomly drawn from different distributions (slow and fast contexts, bottom). At some point, there is an uncued transition to a novel distribution, consisting of a single input interval, ranging between the means of the slow and fast contexts. Center. Speed of neural activity in the RNN vs the projection of neural activity onto the estimated context dimension during the measurement epoch (see Methods). Right. Same analysis in the neural data. In the novel uncued context (grey dot), the speed and projection onto context are adjusted to values between the slow and fast contexts.

A direct consequence of this mechanism is that the network becomes sensitive to local variations in the statistics of the intervals within a given context, and adapts its output accordingly. In particular, the network’s output on a given trial is biased toward the input interval of the previous trial (Fig. 7 B). Strikingly, we found a similar tendency in monkeys trained on a single context (Fig. 7 B right): animals respond generally faster following a short interval, and slower following a long interval. Note that this history effect is abolished in the networks when the updating of the control input is turned off (Fig. 7 B middle), indicating that this effect specifically emerges from contextual input adjustments.

These behavioral results suggest that animals may rely on an updating mechanism similar to that implemented in the networks to remain tuned to changing stimulus statistics. To test the predictions of the model directly at the level of neural dynamics, we developed a novel experiment that combined context-based and adaptation-based timing. Each session was divided in two parts. In the first part, animals reproduced measured intervals under the same conditions as previously, i.e., they were exposed to two alternating interval distributions explicitly cued via the color of the fixation spot. In the second part of the experiment, following an uncued switch, the distribution corresponding to one of the two contexts was covertly changed to an intermediate distribution, while the color of the fixation point remained unchanged (Fig. 7 C left). The specific predictions of our model were that: (i) the level of the pre-stimulus contextual input adapts to an intermediate value along the identified contextual axis; (ii) the speed of the dynamics following the Ready pulse adjusts according to the level of the contextual input.

We analyzed DMFC population activity in this task, and compared it to the predictions of our low-rank networks, similar to Fig. 6 C left. As expected, in the RNNs, after the switch, the network learned the new value of the contextual input, and this value was associated with an intermediate value of speed (Fig. 7 C, grey dots). The same key findings were observed in the neural data: we projected population activity post-switch along the context dimension defined pre-switch, and found that this projection as well as the speed of dynamics were adjusted to intermediate values appropriate for the new distribution (Fig. 7 C, right). This result is particularly notable, since it demonstrates a non-trivial match between neural data and our low-rank networks constrained to adapt via changes in a tonic input that encodes context.

Overall, these analyses provide compelling evidence that a low-dimensional input control strategy provides an efficient way to promote both generalization and adaptability for flexible timing, and is likely at play in the brain.

## 3 Discussion

Examining recurrent neural networks trained on a set of flexible timing tasks, in this study we show that controlling low-dimensional dynamics with tonic inputs enables a smooth extrapolation to inputs and outputs well beyond the training range. Reverse-engineering and theoretical analyses of the recurrent networks demonstrated that the underlying mechanism for generalization relied on a specific geometry of collective dynamics. Within a given condition, collective dynamics evolved along non-linear manifolds, while across conditions, tonic cues modulated the manifolds along an orthogonal direction. This modulation parametrically controlled the speed of dynamics on the manifolds while leaving their geometry largely invariant. We demonstrated that this mechanism leads to fast adapting responses in changing environments by adjusting the amplitude of the tonic input, while reusing the same recurrent local network. Population analyses of neural activity recorded while monkeys adaptively solved a time-interval reproduction tasks confirmed the key geometric and dynamic signatures of this mechanism.

At the algorithmic level, the RNNs that generalized to novel stimuli suggested a computational principle for dynamical tasks that makes a clear separation between recurrent dynamics and two types of inputs based on their function and timescale. First, the scaffold for time-varying dynamics is crafted by recurrent interactions that generate manifolds in neural state-space. Second, on the level of individual trials, fast inputs place neural trajectories at the suitable locations in state-space that allow time-varying responses to unfold. Third, tonic inputs, which are constant during the trial duration, provide the parametric control of the dynamics on the manifold by shifting trajectories to different regions of state-space. These tonic inputs can however vary at the timescale of trials, and encode the contextual information necessary to adapt to the changing statistics of the environment (Meirhaeghe et al., 2021). We expect that this principle of a separation of inputs along different behaviorally-relevant timescales extends to other parametric forms of control (Lake et al., 2017; Hardy et al., 2018; Rajalingham et al., 2021), and provide a fundamental building block for neural networks that implement more complex internal models of the external world.

A key objective of computational modeling is to make falsifiable predictions in experimental recordings; our work on RNNs supplies predictions for both generalization and adaptation. We performed experiments on primates that directly test some of the predictions on adaptation. We found a behavioral bias towards previous trials, compatible in our model with a predictive error signal required for adaptation. At the level of neural recordings, in line with the modeled tonic input controlling neural speed, we found that neural trajectories in dorsomedial frontal cortex adaptively varied along a contextual axis that determined the speed of neural dynamics. On the other hand, we did not have the data for validating our predictions on generalization (i.e., zero-shot learning). Nevertheless, based on the adaptation results, we expect that when animals generalize, their neural activity adjusts the tonic input within a single trial. This change in the input to the local network could be provided either through an external input present during the whole trial duration (e.g., an explicit instruction, or context cue as in Sohn et al. (2019)), or some transient input that is mapped onto a tonic input in the brain (e.g., in reversal learning paradigms, where the contextual input reflects the inference of sudden switches in the environment (Izquierdo et al., 2017; Sarafyazd and Jazayeri, 2019)).

We have shown that low-dimensional recurrent dynamics can be flexibly controlled by a tonic input that varies along one dimension. It remains an open question how low-dimensional the dynamics needs to be in order to be controllable with an external tonic input. In our case, we enforced the minimal dimensionality required to solved the task (up to three dimensions for the MWG task), and showed that networks generalize when provided with a tonic input. In the other extreme, trained networks without any dimensionality constraint did not generalize with a tonic input. These results suggest that flexible cognitive tasks that require more time to be learned are either high-dimensional and require additional mechanisms beyond a tonic input for generalization, or that the extra time is needed to reduce the dimensionality of the network dynamics. These insights provide a test bed to develop more efficient learning curricula for RNNs and eventually animal training, not only in terms of generalization, but also learning speed. The minimal dimensionality was achieved by directly constraining the rank of the connectivity matrix in each task (Dubreuil et al., 2022). The low-rank connectivity has the added advantage of providing an interpretable mathematical framework for the analysis of network dynamics (Mastrogiuseppe and Ostojic, 2018; Schuessler et al., 2020a; Beiran et al., 2021). However, it is possible that variations in the implementation of the RNNs, such as specific types of regularization in unconstrained networks (Sussillo et al., 2015) and other details of the learning algorithm or neural architecture may equally constrain neural activity to low-dimensional dimensional dynamics and induce comparable or better generalization. From that point of view, the minimal rank constraint used here can be seen as a particular type of inductive bias (Neyshabur et al., 2015; Sinz et al., 2019; Bordelon and Pehlevan, 2021; Canatar et al., 2021) for temporal tasks.

How and where tonic inputs are originated remains to be elucidated. The electrophysiology data in this study recorded exclusively from dorsomedial frontal cortex, while the RNN model is agnostic to the origin of the tonic inputs. Thalamic activity has been found to be a candidate for the tonic input that flexibly controls cortical dynamics (Rikhye et al., 2018; Wang et al., 2018; Logiaco et al., 2021). Nevertheless, when the tonic input provides contextual information, as in the MWG+Ctxt task, other cortical and subcortical brain areas are likely recruited for providing a signal that integrates statistics from past events (Paton and Buonomano, 2018; Monteiro et al., 2021). Concurrently, contextual modulation of firing rates has been characterized in different areas of prefrontal cortex (Rigotti et al., 2010; Bouchacourt et al., 2020; Meirhaeghe et al., 2021). Whether this contextual information is integrated and broadcast in a localized cortical region, or emerges from distributed interactions of multiple brain areas, will need to be determined by causal perturbation experiments across brain regions.

## Acknowledgements

MB, MJ and SO were supported by the CRCNS project PIND funded through the National Institute of Health (NIMH: 1R01MH122025-01) and French Agence Nationale de la Recherche (ANR-19-NEUC-0001-01). MB was supported by the Ecole de Neurosciences de Paris. NM was supported by a Whitaker Health Sciences Fund Fellowship. HS was supported by a BBRF Young Investigator grant. SO was supported by the program “Ecoles Universitaires de Recherche” ANR-17-EURE-0017. MJ was supported by the Simons Foundation, the McKnight-Endowment Fund for Neuroscience and the McGovern Institute. The authors would like to thank Adrian Valente for discussions and the software library for training recurrent neural networks, developed initially for Dubreuil et al. (2022).

## Code availability

Code and trained networks will be made available upon publication.

## 4 Methods

### Recurrent neural network dynamics

We trained recurrent neural networks (RNNs) consisting of *N* = 1000 units with dynamics given by Eq. (1). We simulated the network dynamics by applying Euler’s method with a discrete time step Δ*t*. The noise source *η*_*i*_ (*t*) was generated by drawing values from a zero-mean Gaussian distribution at every time step.

The readout of the network was defined as

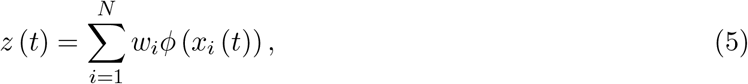

a linear combination of the firing rates of all network units, along the vector ***w*** = {*w*_*i*_}_*i*=1…*N*_.

We considered networks with constrained low-rank connectivity as well as unconstrained networks. In networks of constrained rank *R*, the connectivity matrix was defined as the sum of *R* rank-one matrices

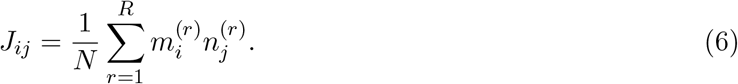

We refer to vectors 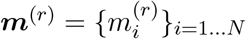 and 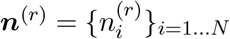 as the *r*−th left and right connectivity vectors for *r* = 1 … *R*.

#### Training

Networks were trained using backpropagation-through-time (Werbos, 1990) to minimize the loss function defined by the squared difference between the readout *z*_*q*_ (*t*) of the network on trial *q* and the target output 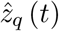 for that trial. The loss function was written as

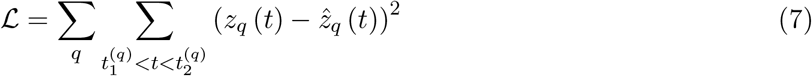

where *q* runs over different trials, and 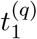 and 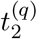 correspond to the time boundaries taken into account for computing the loss function (specified in task definitions below).

The network parameters trained by the algorithm were the components of input vectors ***I***^**(*s*)**^, the readout vector ***w***, the initial network state at the beginning of each trial ***x*** (*t* = 0), and the connectivity. In networks with constrained rank *R*, we directly trained the components of the connectivity vectors ***m***^(*r*)^ and ***n***^(*r*)^, i.e. a total of 2*R* × *N* parameters. In networks with unconstrained rank, the *N* ^2^ connectivity strengths *J*_*ij*_ were trained.

We used 500 trials for each training set, and 100 trials for each test set. Following Dubreuil et al. (2022), we used the ADAM optimizer (Kingma and Ba, 2015) in pytorch (Paszke et al., 2017) with decay rates of the first and second moments of 0.9 and 0.999, and learning rates varying between 10^−4^ and 10^−2^. The remaining parameters for training RNNs are listed in Table 1. The training code was based on the software library developed for training low-rank networks in Dubreuil et al. (2022).

**Table 1:** Parameters for trained RNNs

|  |  |
| --- | --- |
| number of units $N$ | 1000 |
| single-unit $\tau$ | 100 ms |
| standard deviation $\eta_i$ | 0.08 |
| integration step $\Delta t$ | 10 ms |
| initial $g_0$ | 0.8 |
| initial $\sigma_0$ | 0.8 |

In networks with constrained rank, we initialized the connectivity vectors using random Gaussian variables of unit variance and zero-mean. The covariance between components of different connectivity vectors at the beginning of training was defined as

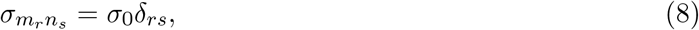

where *σ*_0_ = 0.8 and *δ*_*rs*_ is the Kronecker delta function. These initial covariances between connectivity vectors generated collective activity effectively that was slower than the membrane time constant, which was useful to propagate errors back in time during learning (Schuessler et al., 2020b). In networks where the rank was not constrained, the initial connectivity strengths *J*_*ij*_ were drawn from a Gaussian distribution with zero mean and variance *g*_0_*/N* ^2^. This initial random connectivity is full rank, which leads to connectivity matrices after training that are also full rank (Schuessler et al., 2020b). The input and readout vectors were initialized as random vectors with mean zero and unit variance, randomly correlated with the connectivity vectors.

In the MWG task with contextual cue, networks were initialized using solutions from RNNs trained on the MWG task for initialization. We implemented a two-step method for training RNNs on the MWG+Ctxt task. First, only input and output weights were trained. In a second step, we trained both inputs and output together with the recurrent weights.

In networks with constrained rank, the rank *R* was treated as a hyperparameter of the model. We trained networks with increasing fixed rank, starting from *R* = 1 (see Fig. S1). The minimal rank is defined as the lowest rank *R* for which the loss is comparable to the loss after training a full-rank network (Dubreuil et al., 2022).

### Flexible timing tasks for RNNs

We considered three flexible timing tasks, Cue-Set-Go (CSG, Wang et al. (2018)), Measure-Wait-Go (MWG) and Measure-Wait-Go with context (MWG+Ctxt). All tasks required producing a time interval *t*_*out*_ after a brief input pulse which we denote as ‘Set’. In the three tasks, the input pulse ‘Set’ was defined as an instantaneous pulse along the vector ***I***_*set*_, and indicated the beginning of the output time interval. The target output interval *t*_*out*_ depended on other inputs given to the network, specific to each task and detailed below. The target output in the loss function was designed as a linear ramp (Wang et al., 2018), that started at value −0.5 when the ‘Set’ signal is received, and grew until the threshold value +0.5 (Fig. 1 A-B, Fig. 2 A). The output interval *t*_*out*_ was defined as the time elapsed from the time of ‘Set’ until the time the output reached the threshold value.

The considered time window in the loss function (Eq. 7) included the ramping epoch as well as the 300 ms that preceded and followed the ramp, where the target output was clamped to the initial and final values. For training, we used four target intervals ranging between 800 ms and 1550 ms, about one order of magnitude longer than the membrane time constant of single units. In a small fraction of trials, *p* = 0.1, we omitted the ‘Set’ signal. In that case, the target output of the network remained at the initial value −0.5.

#### Cue-Set-Go task

The target output interval *t*_*out*_ was indicated by the amplitude of a ‘Cue’ input presented before the ‘Set’ signal. The ‘Cue’ input was constant for each trial and present throughout the whole trial duration, along the spatial vector ***I***_*cue*_. In each trial, the ‘Set’ signal was presented at a random time ranging between 400 and 800 ms after trial onset. For training, we used four different cue amplitudes ranging from 0, corresponding to the shortest interval, to 0.25, for the longest interval.

#### Measure-Wait-Go task

The target output interval *t*_*out*_ was indicated by the temporal interval between two pulse inputs along vectors ***I***_1_ and ***I***_2_. Following a random delay ranging between 200 and 1500 ms after the second input, the ‘Set’ input indicated the beginning of the production epoch. Four different input intervals, ranging from 800 ms to 1550 ms were used for training.

#### Measure-Wait-Go with context

We added to the MWG task a tonic contextual input along ***I***_*ctxt*_, that was present during the whole trial duration and covaried with the average duration of the target interval. For this variant of the task, we used eight target intervals for training. Four of them ranged between 800 ms and 1550 ms. In trials with those target intervals, the amplitude of the contextual input was *u*_*ctxt*_ = 0. The other four ranged between 1600 ms and 3100 ms, and the associated amplitude of the contextual input was *u*_*ctxt*_ = 0.1.

In all tasks the first input pulse was fed to the network at a random point in time between 100 and 600 ms after the beginning of each trial.

### Performance measure in timing tasks

We summarized the performance in a timing task by the output time interval *t*_*out*_ generated by the network in each trial. Networks trained to produce a ramping output from −0.5 to 0.5 do not always reach exactly the target endpoint 0.5, but stay close to this value. To avoid inaccuracies due to this variability, we estimated the produced time interval by setting a threshold slightly lower than the endpoint, at a value of *v* = 0.3. We determined the time *t*_*v*_ elapsed between the ‘Set’ input and the threshold crossing. The produced time interval in each trial was then estimated as *t*_*out*_ = *t*_*v*_*/v*. In trained networks where the readout activity did not reach the threshold (e.g., Fig. S3), the threshold crossing was estimated as the time in which the readout was the closest value to threshold.

### Dimensionality of neural activity

For trained networks, we assessed dimensionality (Fig. 2 B) using two complementary approaches: the variance explained by the first principal components, and the participation ratio. For the variance explained, we focused on the production epoch, common to all tasks, and defined as the time window between the ‘Set’ pulse and the threshold crossing. We subsampled the time points for each trial condition so that each of them contributes with the same number of time points. For the participation ratio (Fig. 2 B inset), we focused on the whole trial duration, to better show the differences in dimensionality between the different tasks and connectivity constraints.

We applied principal component analysis to the firing rates *ϕ*(*x*_*i*_ (*t*)) of the recurrent units for every different trial in a given task. The principal component decomposition quantifies the percentage of variance in the neural signal explained along orthogonal patterns of network activity. Due to the presence of single-unit noise in the RNNs, all principal components explain a fixed fraction of variance in the neural signal. The dimensionality can be defined in practice as the number of principal components necessary to account for a given percentage of the neural signal. Alternatively, the dimensionality of two different RNNs can be compared by comparing the distribution of explained variance across the first principal components as shown in Fig. 2 B.

Additionally, we quantified the dimensionality by means of the *participation ratio* (Abbott et al., 2011; Litwin-Kumar et al., 2017; Susman et al., 2021), defined as:

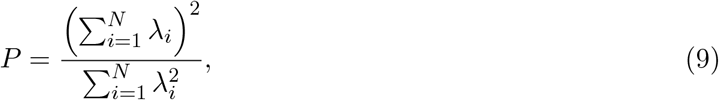

where *λ*_*i*_ correspond to the eigenvalues of the covariance matrix of the neural signal *ϕ*(*x*_*i*_ (*t*)),

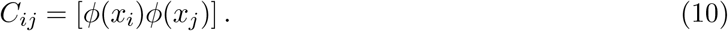

The square brackets denote the time-average and trial-average over the time window of interest. The participation ratio is an index that not only takes into account the dimensionality of neural trajectories, but also weighs each dimension by the fraction of signal variance explained.

### Analysis of trained RNNs

For the analysis of trained networks, the dynamics of RNNs with constrained low-rank were reduced to a low-dimensional dynamical system (Eq. 4), as detailed here. For any rank-*R* RNN, the connectivity matrix ***J*** can be decomposed uniquely using singular value decomposition as the sum of *R* rank-one terms (Eq. 6) where the left (resp. right) connectivity vectors {***m***^(*r*)^} _*r*=1…*R*_ (resp. {***n***^(*r*)^} _*r*=1…*R*_) are orthogonal to each other.

The dynamics of the network (Eq. 1), written in vector notation, read:

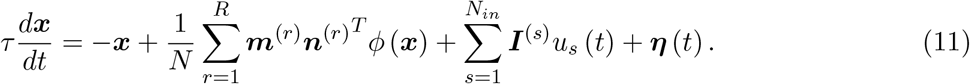

We decompose each input vector ***I***^(*s*)^ into the orthogonal and parallel components to the left connectivity vectors:

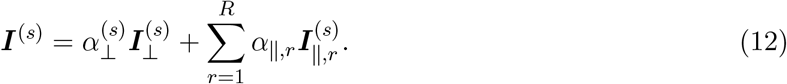

where the constants 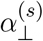 and 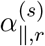 for *r* = 1, …, *R* and *s* = 1, …, *N*_*in*_ indicate the fraction of the input pattern that correspond to each basis vector.

The vector of collective activity (that represents the total input received by each unit) ***x*** (*t*) was then expressed in the basis given by the left connectivity vectors {***m***^(*r*)^}_*r*=1…*R*_ and the orthogonal input vectors components 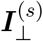 (Beiran et al., 2021; Dubreuil et al., 2022):

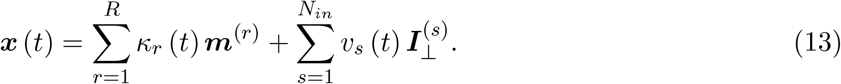

The time-dependent variables ***κ*** = {*κ*_*r*_}_*r*=1…*R*_ represent the projection of the activity along the *recurrent subspace* spanned by the recurrent connectivity vectors {***m***^(*r*)^}_*r*=1…*R*_, and 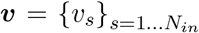 represent the projection of the activity along the *input-driven subspace*. Altogether, we refer to subspace spanned by the left connectivity vectors and orthogonal inputs as the *embedding subspace*. The projection of the activity ***x*** (*t*) along the connectivity vector ***m***^(*r*)^ was in practice calculated as:

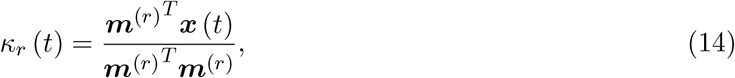

and similarly for the input variables.

Inserting Eq. (13) in Eq. (11), and separating each term along orthogonal vectors of the embedding space, we obtain a set of differential equations for the recurrent and input-driven variables:

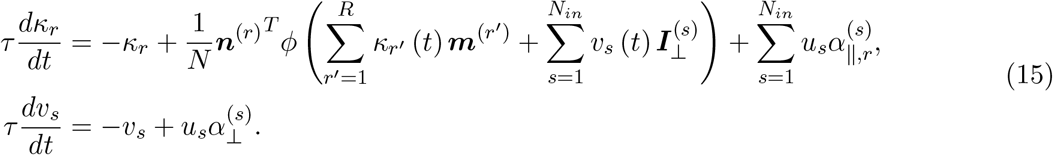

This analysis effectively reduces a high-dimensional dynamical system (Eq. 11, *N* variables) to a lower dimensional dynamical system (Eq. 15, *R* + *N*_*in*_ variables) based on the fact that the recurrent connectivity is rank *R*.

Note that the input-driven variables ***v*** are a temporally filtered version of the input variables ***u*** at the single unit time constant *τ* (Eq. 15). Therefore, pulse-like inputs produce a change in the recurrent variables ***κ*** at the timescale given by *τ*. For constant inputs *u****I*** with variable amplitude *u* from trial to trial, as the cue in the CSG task and context in the MWG+Ctxt task, the effect on the dynamics is twofold. First, varying the amplitude is equivalent to shifting the location of the recurrent subspace to a parallel plane in the embedding subspace, because the input-driven variable *v* is different for each trial. Secondly, the dynamics of the recurrent variables are also affected by changes in the amplitude. They read:

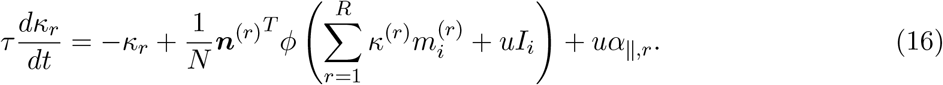

To study the dynamical landscape of low-dimensional activity, we define the speed *q* (Sussillo and Barak, 2013) at a given neural state as a scalar function

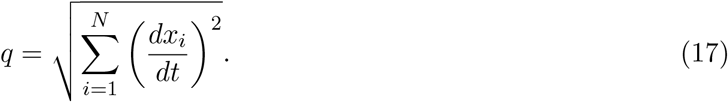

The speed *q* indicates how fast trajectories evolve at a given point in state space. States ***κ*** where the speed is zero correspond to fixed points of the RNN.

In full-rank networks, it is a priori not possible to fully describe the trajectories using only a few collective variables. The speed of the dynamics can however still be calculated as in Eq. (17).

### Non-linear manifolds

A useful approach to analyze the dynamics of low-rank RNNs is to initialize the network at arbitrary initial conditions and visualize the dynamics of the variables ***κ*** in the recurrent subspace, and as a function of time (Fig. S9). We found that before reaching a stable state trajectories with random initial conditions in trained networks appear to converge to non-linear regions of the recurrent subspace, that we refer to as *neural manifolds*.

We therefore devised methods to identify these non-linear manifolds: one exact method, that we used in practice for rank-two networks, and an approximate method, used for rank *R >* 2. The first method consists of initializing trajectories close to all saddle points of the dynamics. In rank-two networks trained on the CSG task, for instance, there are two saddle points, and initializing two trajectories nearby the two opposite saddle points led to a closed curve to which random trajectories converge (solid red line, Fig. S9 A). The manifolds obtained through this method are closely related to the concept of heteroclinic orbits (Rabinovich et al., 2008b,a).

In rank-three networks trained on the MWG task, randomly initialized trajectories converged to a sphere-like manifold (Fig. S3 A). Determining the manifold starting from non-trivial saddle points would require computing a large number of trajectories, to sample all the possible trajectories on the surface of the sphere-like manifold. We instead used an approximate method for determining the manifold. This method consisted in sampling each radial direction of the recurrent subspace, parametrized by the polar and azimuth angles *θ* and *ϕ*, and localizing the non-zero radial distance where the dynamics had minimal speed as defined in Eq. (17) (grey surface, Fig. S9 A; in practice we set a threshold for a minimum distance). Once the manifold was identified, the dynamical landscape on its surface was calculated by locally projecting in two dimensions the vector field on the plane perpendicular to each manifold state. In rank-two networks, the approximate method to determine the manifold led to a curve (red dashed line, Fig. S9 A) which was close to the manifold determined by the first method, in particular at the fixed points, and on the portions of the manifold where the dynamics were slow.

To confirm that the identified manifolds correspond to slow manifolds of the dynamics, we computed the distance of randomly initialized trajectories to the manifold. We found that the distance to the manifold decayed to zero at a timescale given by the membrane time constant (Fig. S9 C). In contrast, projecting the recurrent variables onto a linear readout, we find that trajectories took a much longer time to converge to a stable fixed point of the dynamics (Fig. S9 B).

Networks of rank *R* do not necessarily generate manifolds. As a counter-example, Fig. S9 D-F display the dynamics of a rank-two network, that led to only two non-trivial stable fixed point, and no saddle points. In this case, initializing trajectories randomly led to trajectories that approach the stable fixed points along different curves. The dynamics of the projected activity along the readout in this case converged quickly to the stable fixed points.

### Mean-field low-rank networks

Dynamics in low-rank networks become mathematically tractable in the limit of large networks when the connectivity components of every unit are randomly drawn from a multivariate probability distribution. Here we assumed that the components of connectivity and input vectors were drawn from a zero-mean multivariate Gaussian (Mastrogiuseppe and Ostojic, 2018; Schuessler et al., 2020a; Beiran et al., 2021)

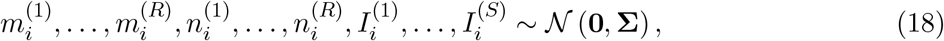

where **Σ** is the (2*R* + *N*_*in*_) × (2*R* + *N*_*in*_) covariance matrix. We introduce the notation *P* (*<u>m</u>, <u>n</u>, <u>I</u>*) = *P*(*m*^(1)^,…,*m*^(*R*),^*n*^(1)^,…,*n*^(R^),*I*^(1)^,…,*I*^(*s*)^ to refer to the joint probability distribution of vector components. Without loss of generality, we fixed the variance of the right connectivity vectors and input vectors to unity; 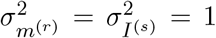. We further assumed that the external input vectors are orthogonal to the left connectivity vectors ***m***^(*r*)^.

The overlap between two vectors ***x*** and ***y*** was defined as their empirical covariance:

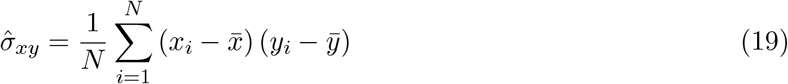

Here 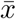 indicates the average value of the components, set to zero in mean-field networks, so that the overlap between two vectors was equivalent to their scalar product. In the large *N* limit, the overlap between vectors ***x*** and ***y*** therefore converges to the covariance *σ*_*xy*_ between their components. Importantly, as the full covariance matrix **Σ** needs to be positive-definite, not all its elements *σ*_*xy*_ are free parameters.

The parameters that determine the dynamics are then the covariances between left and right connectivity vectors and the covariances between input and left connectivity vectors, as we detail below (see also Beiran et al. (2021)). For that reason, we defined the overlap matrix ***σ***_***mn***_ as the matrix with elements 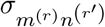, for *r, r*′ = 1, …, *R*, and the covariance vector ***σ***_***In***_ between input and right connectivity vectors as the vector with components 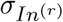. The overlap matrix ***σ***_***mn***_ and vector ***σ***_***In***_ correspond to different subsets of elements of the full covariance matrix **Σ**.

Given these definitions and assuming just one constant input *v***I**, in the limit of large networks *N* → ∞, the sum over *N* units in Eq. (15) can be replaced by the expected value over the Gaussian distribution of connectivity and input vectors. The dynamics of the recurrent variables then read:

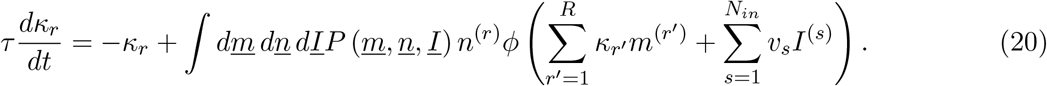

The Gaussian integral in Eq. (20) can be further expressed in terms of the covariances of the probability distribution as (Schuessler et al., 2020a; Beiran et al., 2021; Dubreuil et al., 2022)

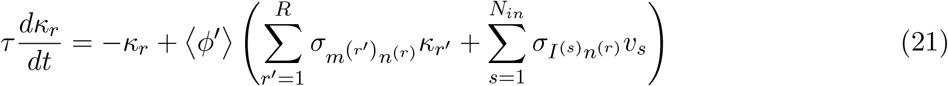

where the state-dependent gain factor ⟨*ϕ*′⟩ was defined as

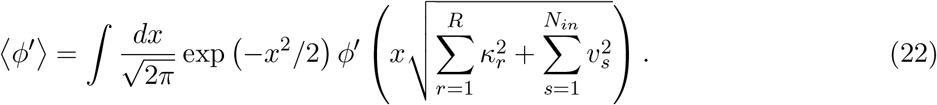

The dynamics plotted in Fig. S10 were generated directly from Eq. (21).

For constant inputs (as in Fig. 5) we define *non-specific inputs* as inputs that are uncorrelated with the connectivity vectors, so that 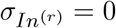 for *r* = 1, …, *R. Subspace-specific inputs* are defined as inputs that are correlated with at least one of the right connectivity vectors 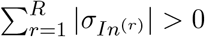.

### Analysis of dynamics in mean-field low-rank networks

We first focus on fixed points of the dynamics in Eq. (21) in absence of inputs. The fixed points correspond to values of *κ*_*r*_ for which the r.h.s of Eq. (21) is zero, which leads to:

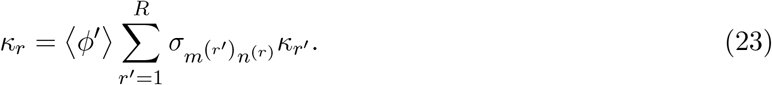

Since the gain (*ϕ*′) is implicitly a function of *κ*_*r*_ (Eq. 22), Eq. (23) is a set of *R* non-linear equations, the solutions of which depend only on the overlap matrix ***σ***_***mn***_. Mathematical analyses show that non-zero fixed points correspond to states along the eigenvector directions of the overlap matrix ***σ***_***mn***_***κ***_*r*_ = *λ*_*r*_***κ***_*r*_, for eigenvectors whose eigenvalues are larger than one, *λ*_*r*_ *>* 1 (Schuessler et al., 2020a; Beiran et al., 2021). Only the fixed points corresponding to the largest eigenvalue are stable, while all the other non-trivial fixed points are saddle points (Schuessler et al., 2020a; Beiran et al., 2021).

If the overlap matrix ***σ***_***mn***_ has a degenerate eigenvalue *λ*_*r*_, with at least two orthogonal eigenvectors ***κ***_*r*1_ and ***κ***_*r*2_ (as is the case when the overlap matrix ***σ***_***mn***_ is proportional to the identity matrix), each possible linear combination of the two eigenvectors leads to a fixed point. Consequently, the system displays a continuous set of fixed points with identical stability, symmetrically located around the origin. In rank-two networks, such a degenerate overlap matrix leads to a ring attractor (Fig. S10 B) (Mastrogiuseppe and Ostojic, 2018; Beiran et al., 2021; Darshan and Rivkind, 2022). All fixed points on the ring are marginally stable, as the direction tangent to the ring corresponds to a zero eigenvalue. In rank-three networks, such symmetry in the connectivity leads to a spherical attractor.

In finite networks, however, the overlap matrix ***σ***_***mn***_ is perturbed by the finite-size sampling, so that it is never exactly proportional to the identity matrix. Small perturbations of the overlap matrix break the continuous symmetry of the attractor, generally leading to a pair of stable fixed points and saddle points on the former attractor, but keeping the speed of the dynamics slow along its surface. The location of the fixed points are then given by Eq. (23). If the perturbation of the symmetric overlap matrix has orthogonal eigenvectors, two stable fixed points and two saddle points are generated on the manifold along orthogonal directions (Fig. S10 C). In contrast, if the perturbation has non-orthogonal eigenvectors, the stable fixed points and saddle points are arranged closer to each other along the manifold (Fig. S10 D).

We then studied the effects of the dynamics of mean-field low-rank networks receiving a constant input *v***I** (Fig. 5). The fixed point equation with an external input reads:

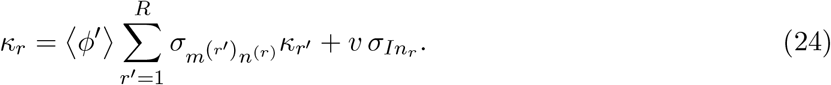

Tonic inputs affect the dynamics of low-rank networks in two different ways. One effect is given by the second term in Eq. (24), which is additive and directed along the vector ***σ***_***In***_. The second effect corresponds to changes in the gain ⟨*ϕ*′⟩ which decreases monotonically towards zero when *v* is increased (Eq. 22). Such changes correspond to the multiplicative factor in Eq. (24), and depend only on the strength of the input, but not its direction. Non-specific tonic inputs lead only to the second effect because for them 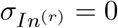 for *r* = 1, …, *R* (Fig. 5 A-B), while subspace-specific inputs combine both effects (Fig. 5 D-E).

### Ready-Set-Go task in monkeys

Details about the behavioral task in monkeys can be found in Sohn et al. (2019) and Meirhaeghe et al. (2021). Briefly, two macaque monkeys (monkey G and H) performed a time interval reproduction task known as the ‘Ready-Set-Go’ (RSG) task (Jazayeri and Shadlen, 2015; Sohn et al., 2019; Meirhaeghe et al., 2021) Each trial began with animals maintaining their gaze on a central fixation point (white circle: diameter 0.5 deg; fixation window: radius 3.5 deg) presented on a black screen. Upon successful fixation, and after a random delay (uniform hazard; mean: 750 ms, min: 500 ms), a peripheral target (white circle: diameter 0.5 deg) was presented 10 degrees left or right of the fixation point and stayed on throughout the trial. After another random delay (uniform hazard; mean: 500 ms, min: 250 ms), the ‘Ready’ and ‘Set’ cues (white annulus: outer diameter 2.2 deg; thickness: 0.1 deg; duration: 100 ms) were sequentially flashed around the fixation point. Following ‘Set’, the animal had to make a proactive saccade (self-initiated Go) toward the peripheral target so that the output interval (*t*_*out*_, between ‘Set’ and ‘Go’) matched the input interval (*t*_*in*_, between ‘Ready’ and ‘Set’). The sample interval, *t*_*s*_, was sampled from one of two discrete uniform distributions, with 5 values each between 480–800 ms for the fast context, and 800–1200 ms for the slow context. The distributions alternated in short blocks of trials (min of 5 trials for G, 3 trials for H, plus a random sample from a geometric distribution with mean 3, capped at 25 trials for G, 20 trials for H), and were indicated to the animal by the color of the fixation point (context cue; red for Fast context, blue for Slow context).

The trial was rewarded if the relative error, |*t*_*out*_ − *t*_*in*_|*/t*_*in*_, was smaller than 0.2. If the trial was rewarded, the color of the target turned green, and the amount of juice delivered decreased linearly with the magnitude of the error. Otherwise, the color of the target turned red, and no juice was delivered. The trial was aborted if the animal broke fixation prematurely before ‘Set’ or did not acquire the target within 3 *t*_*in*_ after ‘Set’. After a fixed inter-trial interval, the fixation point was presented again to indicate the start of a new trial. To compensate for lower expected reward rate in the Long context due to longer duration trials (i.e. longer *t*_*in*_ values), we set the inter-trial intervals of the Short and Long contexts to 1220 ms and 500 ms, respectively.

### Neural data

For Fig. 6, details about the neural recordings related to the RSG task (compared with MWG+Ctxt task) can be found in Sohn et al. (2019) and Meirhaeghe et al. (2021). In summary, the data were obtained from acute recordings in the dorsomedial frontal cortex (DMFC; n=619 neurons in monkey G, n=542 in monkey H). For the neural analysis, the context dimension was defined as follows: we first computed the trial-averaged firing rates (bin size: 20 ms, Gaussian smoothing kernel SD: 40 ms) between ‘Ready’ and ‘Set’ (measurement epoch) for the Short and Long contexts separately. Because the animal did not know beforehand which input interval (*t*_*in*_) was presented on a given trial, we averaged trials irrespective of the *t*_*in*_ value within each context. The context dimension was defined as the unit vector connecting the state at the time of Ready of the Long condition and the state at the time of Ready of the Short condition. We then projected the Ready state of the Short and Long contexts onto the context dimension (Fig. 6 C, bottom left). We normalized the projection such that the mean is zero, and the two average contexts correspond to −0.5 and 0.5. To compute the neural speed in the measurement epoch for each context, we averaged the Euclidean distance between consecutive states between ‘Ready’ and ‘Set’. We used standard bootstrapping (resampling trials with replacement; N=100 repeats) to generate confidence intervals. We normalized the speed such that the maximum average speed is 1.

We performed principal component analysis (PCA) to visualize neural trajectories (Fig. 6 A-B, bottom). PC trajectories were obtained by gathering smoothed firing rates in a 2D data matrix where each column corresponded to a neuron, and each row corresponded to a given time point in the measurement epoch. To obtain a common set of principal components for the two contexts (Short and Long), we concatenated the average firing rates of the recorded neurons for the two contexts along the time dimension. We then applied principal component analysis and projected the original data onto the top 3 PCs, which explained about 75% of total variance. Fig. 6 A was obtained by projecting the trajectories onto the top three principal components. Fig. 6 B was obtained by projecting the trajectories onto the first three principal components and then onto the subspace orthogonal to the estimated contextual axis.

### Geometrical analyses of neural trajectories

To quantify the geometrical analyses of neural trajectories in Fig. 5, we applied the ‘kinematic analysis of neural trajectories’ (KiNeT) framework developed in Remington et al. (2018a). In particular, we used three different geometrical assessments to quantify the three properties observed in the data visualization: (i) low-dimensional modulation of trajectories based on context, (ii) invariance of trajectories along a subspace of neural state-space and (iii) speed modulation of trajectories based on the context signal.

For the low-dimensional modulation of trajectories, we computed the angle Θ between two states in the average trajectory of the slow context and the average trajectory of the fast context. We take as reference trajectory the slow context, and we parameterize the set of successive states by the reference time *t*_*ref*_, the time it takes to reach such state after the first pulse. Secondly, at each state separated by a time lag *τ* from the reference state *t*_*ref*_, we determine which is the closest point in the fast context trajectory, and store this context vector 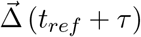 between those two different points. Finally, we compute the angle Θ, in absolute value, between the pairs of context vectors 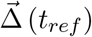 and 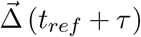. If context perfectly separates neural trajectories along a one-dimensional axis during the measurement epoch, the angle Θ should be zero for all values of time lags *τ* and reference states *t*_*ref*_. Fig. S13 A shows the average angle for all possible reference times as a function of time lag, while Fig. 5 A shows the mean of the average angle for all different time lags. As a control, we compute surrogate context trajectories where we randomly shuffle the context labels of neural trajectories, and compute the angles Θ.

To test the invariance of trajectories along dimensions orthogonal to the context modulation axis (Fig. 5 B), we similarly calculate the distance between the average neural trajectories in each context after removing the projection of neural trajectories along the average context dimension, as calculated in Fig. 5 A. We use a different set of trials for the reference trajectory and the target neural trajectories. As a control, we calculate the distance between trajectories after removing the projection of neural trajectories along a random direction (average over 40 different random directions in state space). If trajectories largely overlap with each other, we expect to find a very short distance between trajectories after removing the context axis. To avoid artifacts due to high-dimensional noise in the recordings, and better compare with the three-dimensional PC visualization, we kept the three first principal components of neural trajectories for this analysis. Fig. S13 B shows the computed distance as a function of the reference time during measurement.

Finally, we can compute the speed of neural trajectories (Fig. 5 C right) by comparing the time elapsed *t*_*comp*_ to reach the closest point to a state in the reference trajectory, compared to how long it took to reach that state in the reference trajectory *t*_*ref*_. If trajectories evolve at the same speed in the reference and the comparison trajectory, *t*_*comp*_ and *t*_*ref*_ should be equal. If trajectories evolve twice faster in the comparison trajectory, *t*_*comp*_ should be half the value of *t*_*ref*_.

### Adaptation experiment in monkeys

For the adaptation experiment (Fig. 7), we collected new data from DMFC in the same animals (n=110 neurons in H, n=129 in monkey G) while they underwent an uncued transition from two alternating contexts (similar to RSG) to a novel intermediate context. The interval distributions used in the two contexts differed from the ones used in the RSG task. For monkey H, the pre-switch distribution was [720;820;920] ms in the slow context, [560;640;720] ms in the fast context, and the post-switch distribution associated with the slow context was reduced to a single interval equal to 720 ms. For monkey G, the pre-switch distribution was [720;820;920;1020] ms in the slow context, [480;560;640;720] ms in the fast context, and the post-switch distribution associated with the slow context was reduced to a single interval equal to 720 ms. The uncued transition occurred randomly during the session, on trial 714 for monkey H, and trial 603 for monkey G. We used 100 trials immediately following the uncued switch to estimate neural data post-switch. Pre-switch data included all trials before the switch, i.e., nearly 100 trials per interval of each context. All other experimental details were identical to the RSG experiment and can be found in Sohn et al. (2019) and Meirhaeghe et al. (2021).

### Dynamical adaptation of context signal

In order to allow the network model to adaptively adjust the value of the context signal based on input interval statistics (Fig. 7), we add an unsupervised predictive loop (Fig. 7). This loop can be split into two parts. First, it estimates the average input interval to predict the input interval in the next trial, by performing a leaky estimation of the input interval over trials, *t*_*pred*_. For that, the network requires one parameter, the update timescale *τ*. The predicted interval is updated at every trial *n* following:

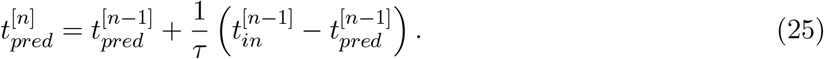

The second term corresponds to the prediction error or mismatch between the input interval and the predicted interval.

Secondly, the module maps the predicted interval to an input amplitude by applying an affine transformation. This mapping requires in general two parameters, the bias term *s*_0_ and the slope *β*, which we assumed have been learned over the course of training. The contextual input amplitude thus reads

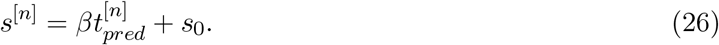

The *β* and *s*_0_ parameters are set so that the mean value of *s* corresponds to the context input values used during training (0 and 0.1 in this case) for the intervals in the slow and fast contexts (1150 ms and 2350 ms, respectively).

### Linear decoders

For Fig. 6 C left and 7 C, we used linear decoders based on cross-validated linear discriminant classification, to readout context from neural activity during the pre-stimulus period in the MWG task with contextual cue. The direction of the classifier is estimated as:

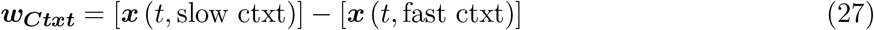

where the square brackets indicate the average across time points and trials of a given context. After the estimation of the decoder, we projected the neural activity of an independent set of trials, different from the training set.

## Supplementary Figures

**Supplementary Figure 1.**
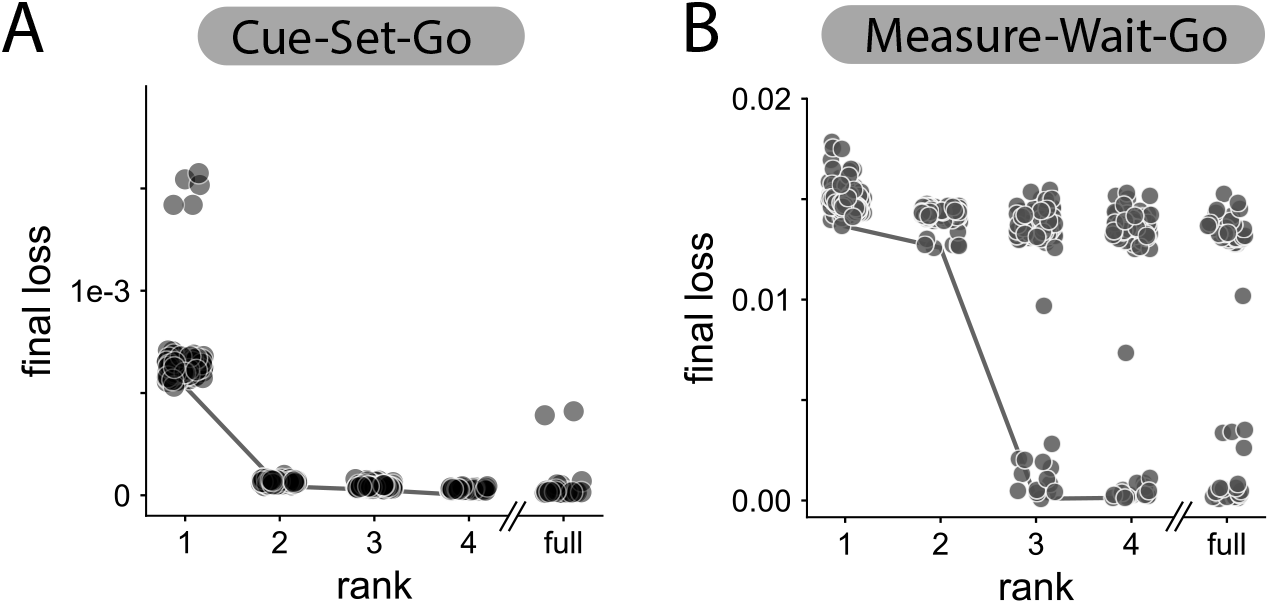
Loss (Eq. 7) of trained low-rank networks as a function of the rank of the connectivity matrix. Dots correspond to 100 different RNNs, the line connects the networks with minimal loss. For comparison, we are showing the loss training full rank networks (rank =*N* =1500). **A** CSG task. The minimal rank for this task is *R* = 2, for which the 100 trained RNNs learned the task. **B** MWG task. The minimal rank is *R* = 3, for which 14 networks learned the task. For rank *R* = 4, 18 networks learned the task. For ranks one and two, no network successfully learned the task.

**Supplementary Figure 2.**
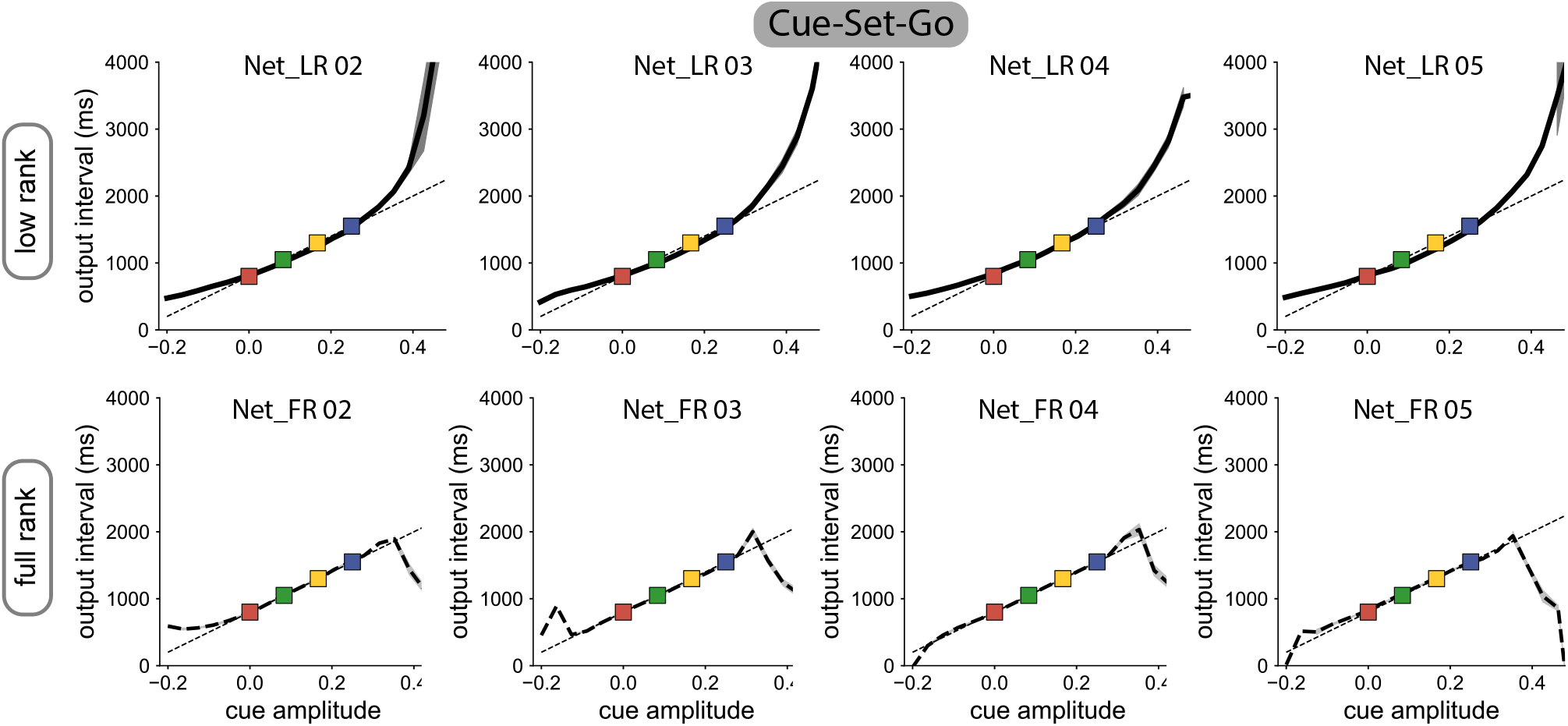
Generalization on Cue-Set-Go task for four different trained RNNs, rank-two (top row) and unconstrained full-rank networks (bottom row), similar to Fig. 1 C. Low-rank networks generalize to higher and lower cue amplitudes, by producing longer and shorter intervals respectively, while unconstrained networks do not.

**Supplementary Figure 3.**
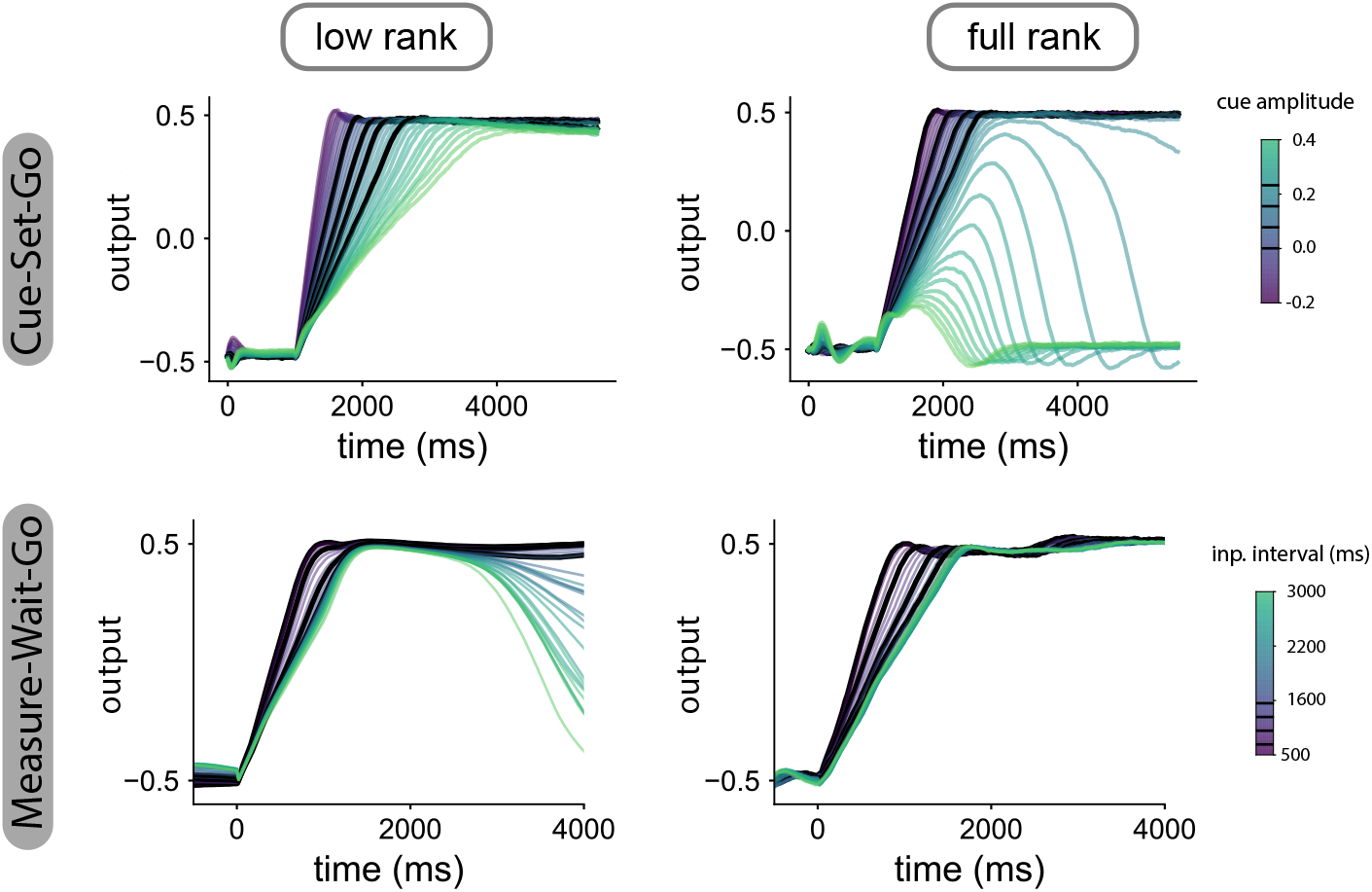
Network output on CSG (top row) and MWG tasks (bottom row) for the networks shown in Fig. 2. Black lines correspond to trials used for training. Only low-rank networks trained on the CSG are able to generalize by modifying the slope of the ramping output (top left). Unconstrained networks trained on CSG do not generate a ramping output for stronger cue amplitudes (top right). In the MWG, both low-rank and unconstrained networks do not produce slower ramps than those used during training (bottom plots).

**Supplementary Figure 4.**
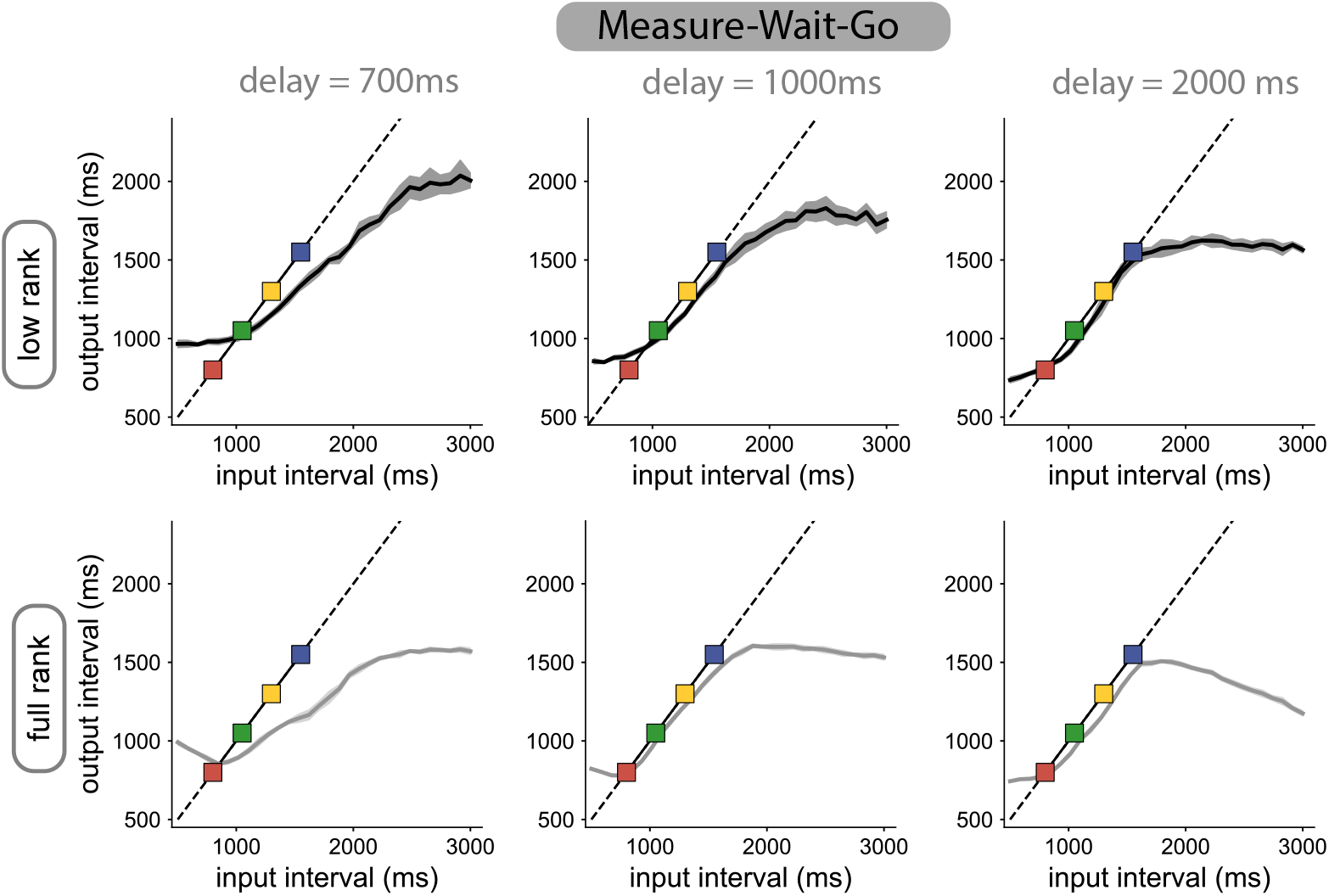
Generalization on the Measure-Wait-Go task, as in Fig. 3, for three different lengths of the delay period. The input-output function is not strongly affected by the duration of the delay period.

**Supplementary Figure 5.**
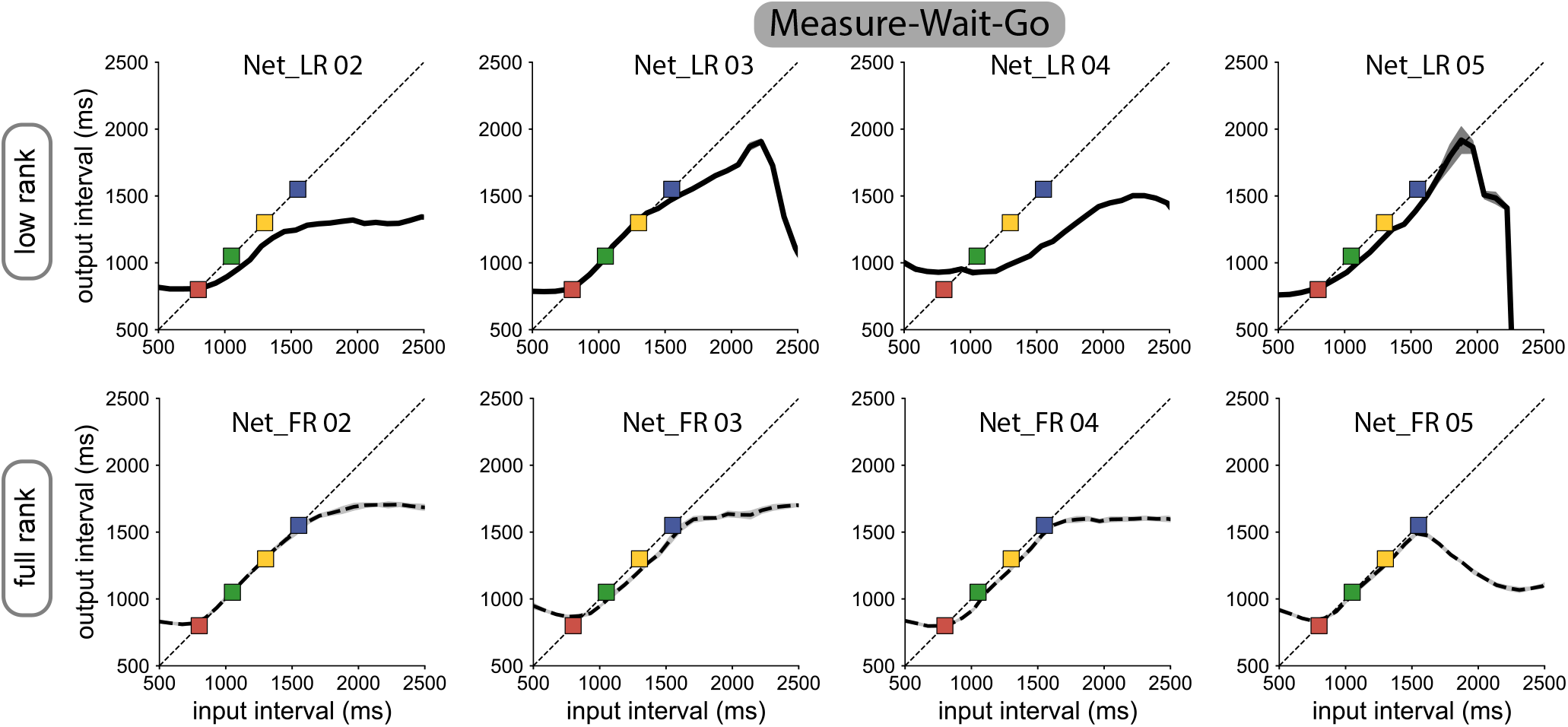
Generalization on the Measure-Wait-Go task for four different trained RNNs, low-rank (top row) and unconstrained (bottom row), similar to Fig. 3. Neither low-rank nor unconstrained RNNs produce intervals much longer than those used in training. **A**

**Supplementary Figure 6.**
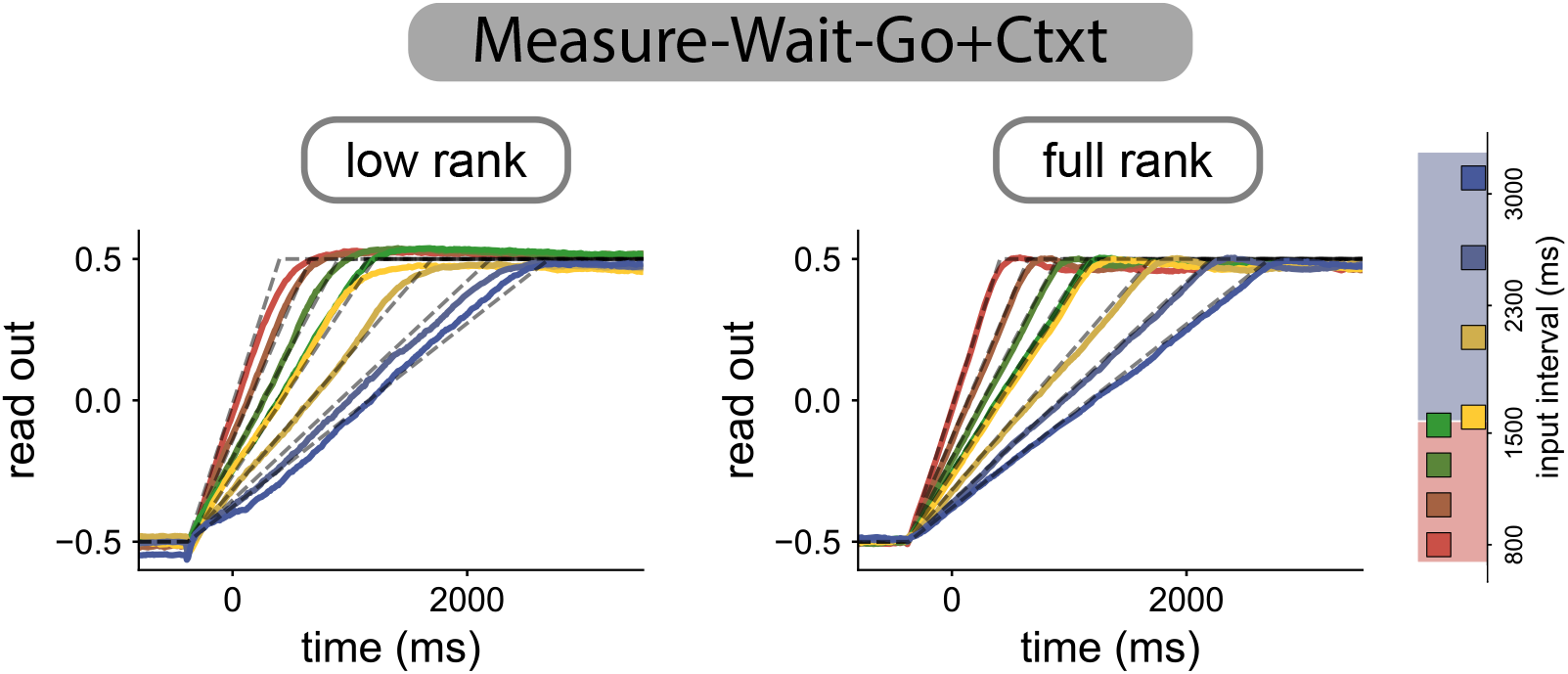
Network output on the MWG+Ctxt task in rank-three (left) and unconstrained (right) RNNs. Both networks are able to produce the eight output intervals used during training, four corresponding to each context (right, red and blue colors).

**Supplementary Figure 7.**
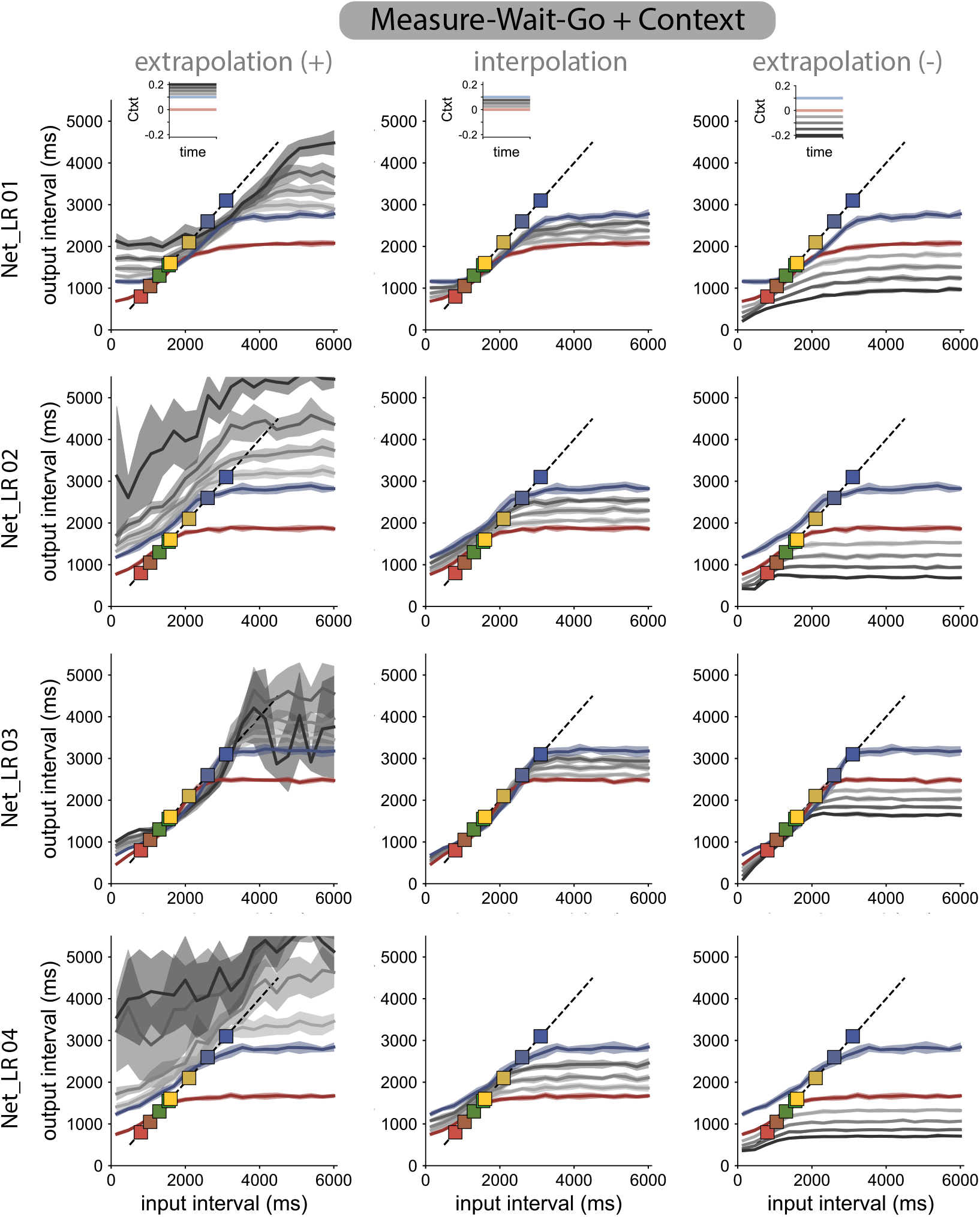
Generalization of four trained rank-three RNNs on the MWG+Ctxt task to contexts not seen during training.

**Supplementary Figure 8.**
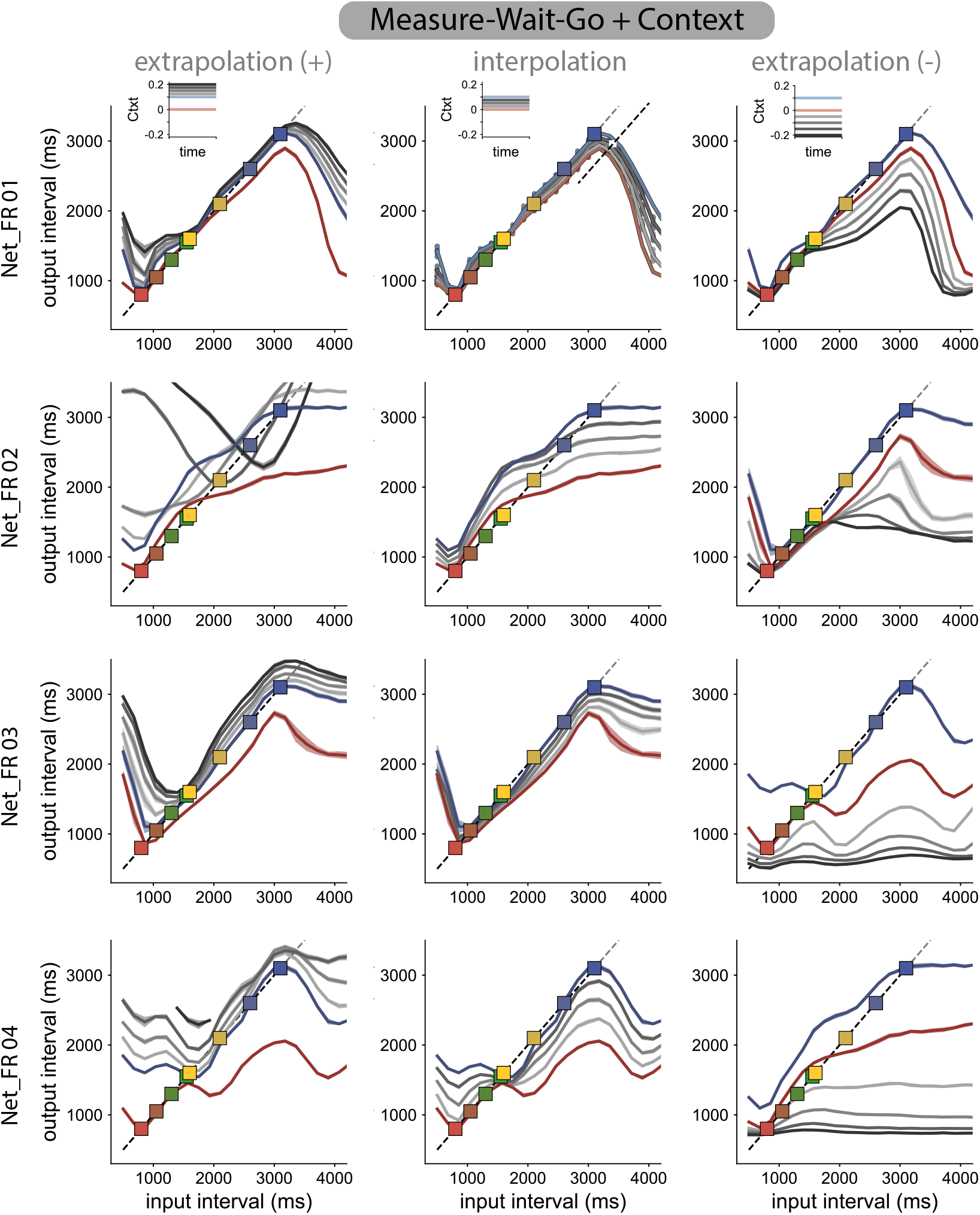
Generalization of four trained full-rank RNNs on the MWG+Ctxt task to novel contexts. Although there is a larger variety of solutions, networks do not generalize smoothly to novel contexts. **A**

**Supplementary Figure 9.**
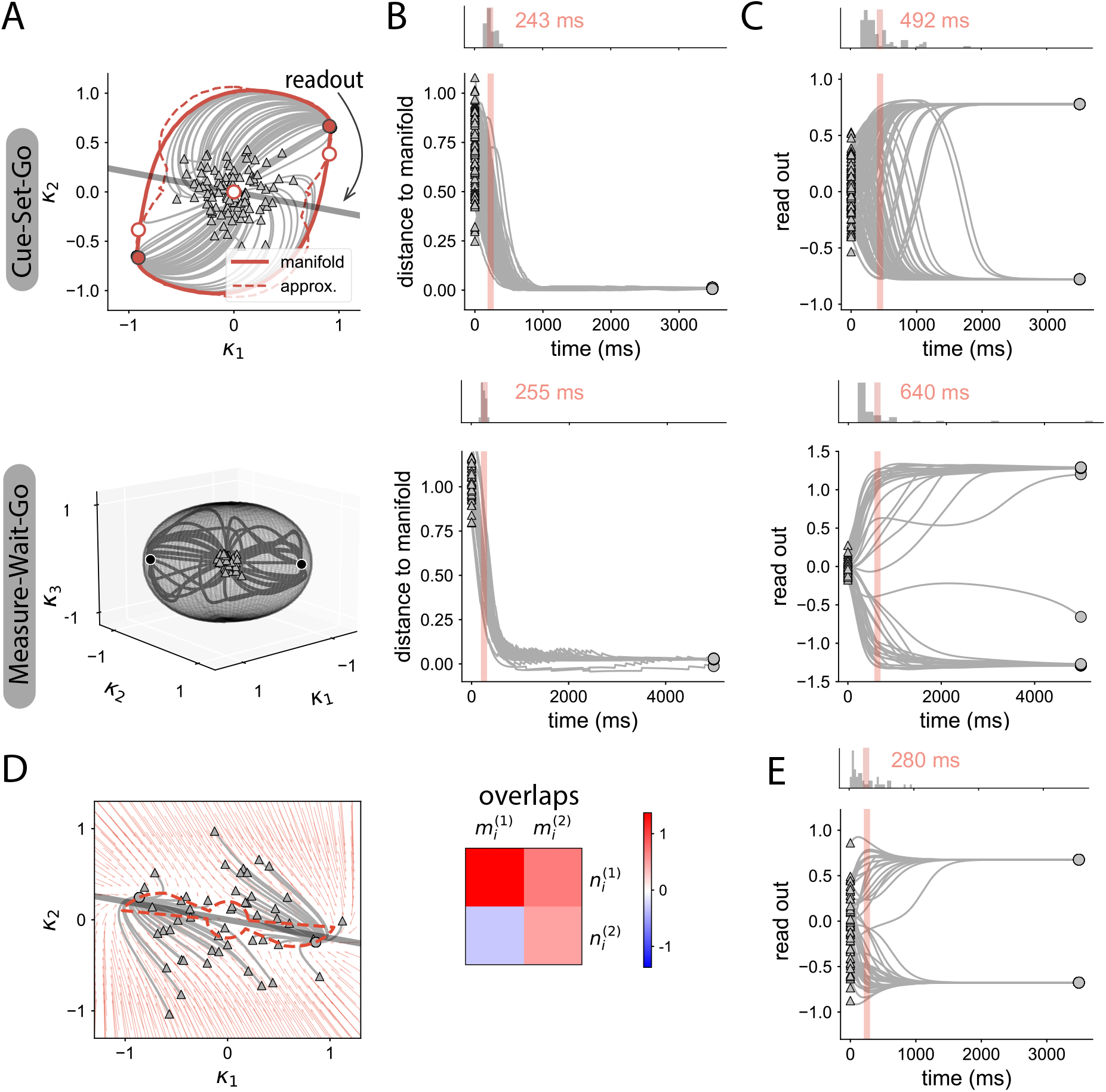
**A** Neural manifolds in low-rank networks trained on the CSG task (top) and MWG task (bottom). Grey lines display trajectories initiated at random states indicated with triangles. White-filled dots are non-stable fixed points, and red/black filled dots are stable fixed points. Top: For the CSG task, the solid red line corresponds to the manifold found by tracking trajectories emanating from saddle points. The red dashed line displays the result of the approximate method for identifying the manifold (see Methods). Bottom: For the MWG task, the spherical-like manifold is shown in grey, determined using the approximate method. **B** Radial distance between randomly initiated trajectories and the manifold. Trajectories converge first to the manifold, and then evolve towards one of the stable fixed points. Red line indicates the average half-time decay from the initial state (see distribution on top inset). **C** Projection of the random trajectories along the readout direction (indicated by grey straight lines in A). Randomly-initialized trajectories converge slowly towards the stable fixed points. **D**-**E** Example of a rank-two network not generating a ring-like manifold. **D** Red arrows correspond to the vector field of the dynamics. Randomly initialized trajectories converge to the non-trivial stable fixed points following different curves. Right: overlap structure between connectivity patterns determining the dynamics. **E** Projection of the random trajectories along the readout direction. Trajectories quickly converge to the non-trivial stable fixed points.

**Supplementary Figure 10.**
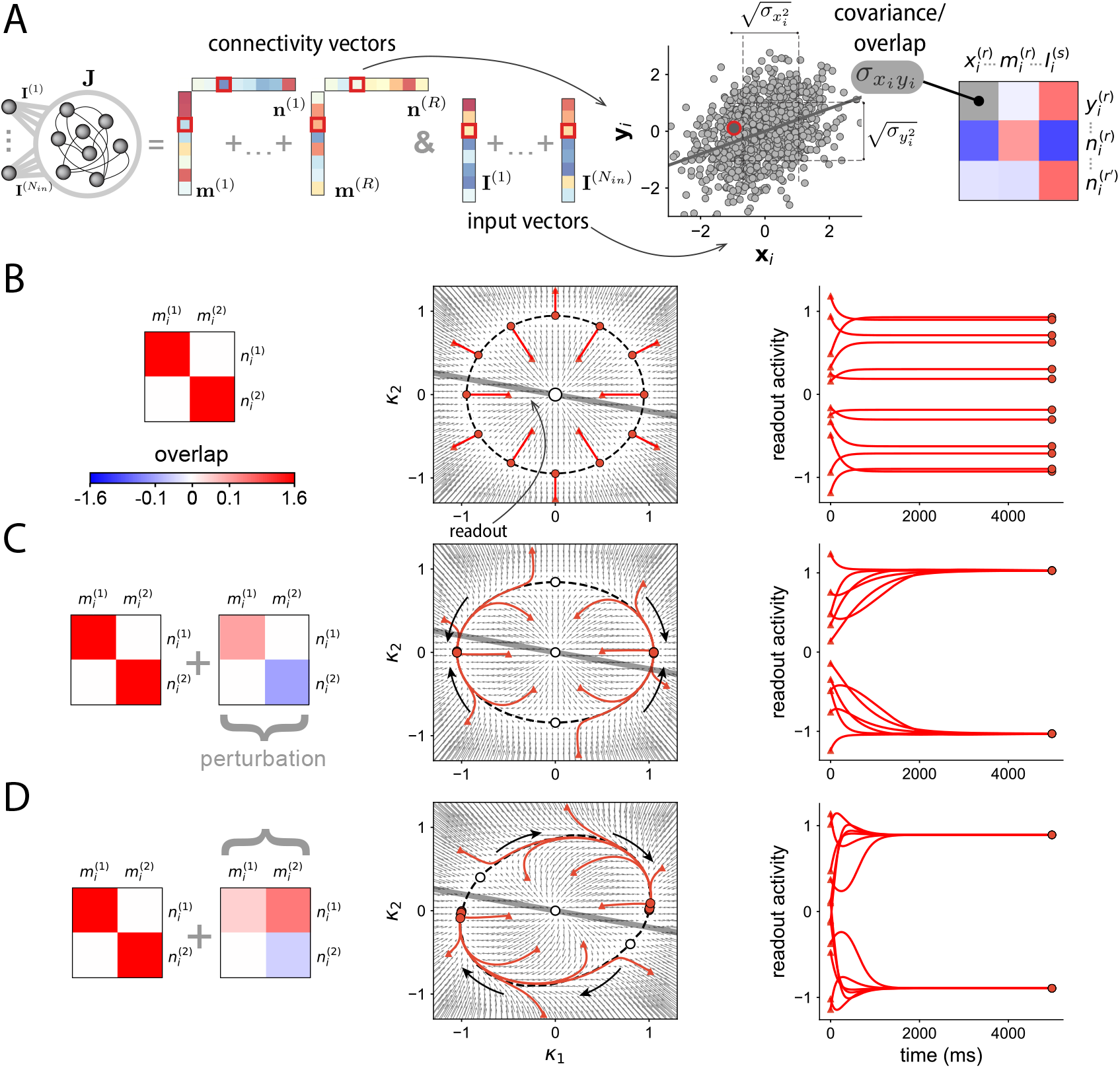
Shaping manifolds by the recurrent connectivity in low-rank networks. **A** Mean-field framework. We assume that the entries of the recurrent connectivity and input vectors are randomly drawn from a zero-mean multivariate Gaussian distribution. The parameters that define the connectivity are therefore the variances 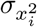 and covariances 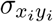, for 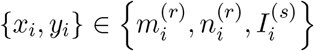, or equivalently the pairwise overlaps between vectors ***m***^(*r*)^, ***n***^(*r*)^, ***I***^(*s*)^. These pairwise overlaps can be represented as entries of a matrix (right) whose values determine the dynamics in the limit of large networks. **B** Ring-attractor dynamics of a rank-two network with overlap structure proportional to the identity (left). Middle: In the mean-field limit, the recurrent connectivity generates a ring attractor in the recurrent subspace. Red lines correspond to trajectories from different initial conditions, that converge towards fixed points. The grey line indicates the readout direction. Right: Temporal trajectories of activity projected along the readout dimension. The trajectories correspond to the mean-field dynamics, given by Eq. (21) (see Methods). **C** Analogous to **B**, in a rank-two network where the overlap structure is perturbed (left, second term). The overlap matrix is still diagonal, but the diagonal entries are different from each other. Middle: This connectivity structure leads to two stable fixed points and two saddle points (white dots) located on the original ring attractor which now forms a slow manifold. Trajectories of activity quickly converge to the ring manifold, and then evolve slowly towards one of the stable fixed points (right). **D** Analogous to **B**, in a rank-two network where the perturbation matrix is not diagonal (left). Middle: The off-diagonal component of the overlap matrix rotates the saddle points closer to the stable fixed points on the slow manifold. Right: The readout activity produces slow ramping signals, as trajectories evolve from one side of the manifold to the other.

**Supplementary Figure 11.**
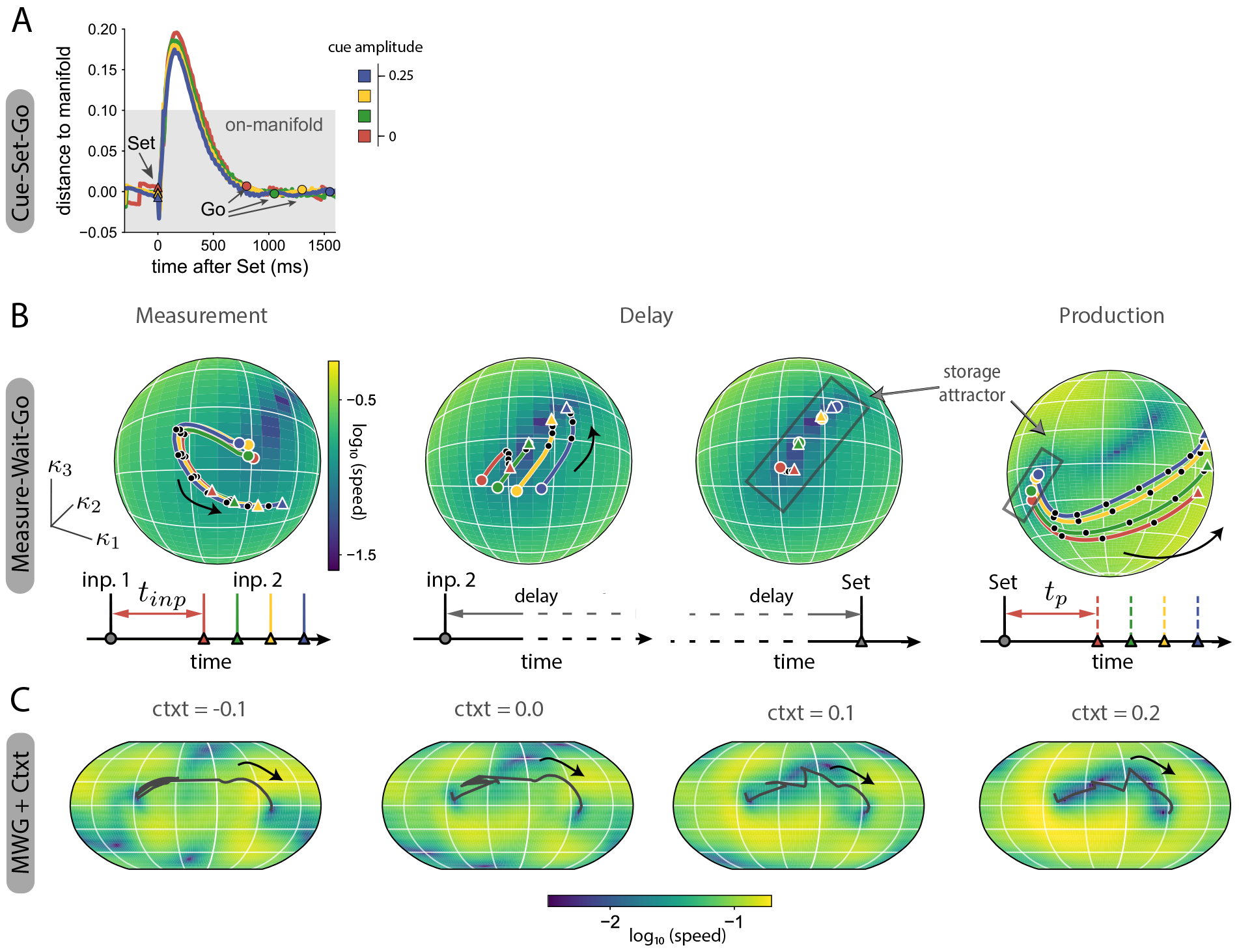
**A** Radial distance to the neural manifold of trajectories in a low-rank network trained the CSG task. The ‘Set’ pulse sends trajectories away from the manifold, but they quickly decay back to it before reaching the final state. **B** Low-dimensional description of neural trajectories in a rank-there network trained on the MWG task. Colored lines are the projections of trajectories on the spherical manifold, corresponding to different input intervals. The colormap on the sphere indicates the speed of the dynamics on the surface of the manifold (blue: slow dynamics, yellow: fast dynamics). Colored dots indicate the state at the beginning of the considered time window, triangles indicate the end of the epoch. Black dots are placed every 200ms. Left: Measurement epoch between the two input pulses. Trajectories are initialized at a non-trivial stable fixed point. The first pulse, indicating the beginning of the measurement epoch, triggers a transient response. The second pulse is received at different states along the evoked transient trajectory, so that the network effectively maps different input intervals to different states on the manifold. Middle panels: activity during the first half (left) and second half (right) of the delay following the second input pulse. Trajectories quickly converge to a slow region of the manifold, where different input intervals are stored in working memory during the remaining delay. The slow region works as a storage attractor: the smallest and largest input intervals stored (red and blue lines) correspond to the extremes of the attractor. Right: Production epoch. The ‘Set’ pulse sends trajectories from the storage attractor towards the other side of the spherical manifold, along parallel trajectories at different speeds. **C** Speed on the surface of the non-linear manifold for different contextual inputs in the rank-three RNN trained on the MWG+Ctxt task. The black lines corresponds to a trajectory solving the task (input interval of 1500 ms). The region explored by the trajectory of the manifold becomes slower as the context input is increased.

**Supplementary Figure 12.**
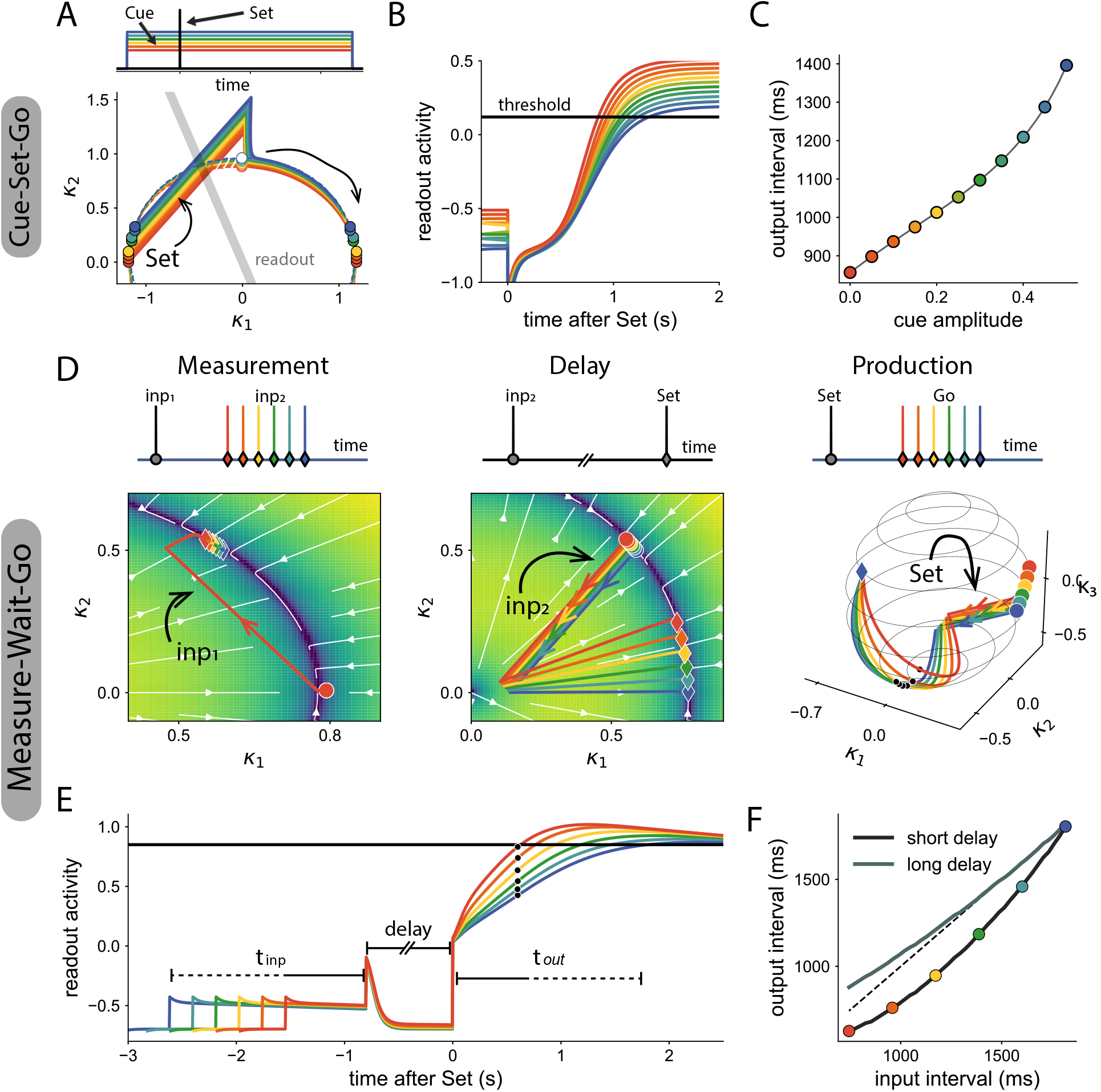
Mean-field low-rank RNNs designed for solving the CSG (**A** - **C**) and MWG (**D** - **F**) tasks. The overlaps between connectivity vectors, and the overlaps of input vectors are chosen by hand to reproduce the mechanisms observed in trained networks. **A** Trajectories solving the CSG task in a rank-two network. The amplitude of the input cue modulates the speed on an invariant manifold (colored dashed lines). Trajectories are initialized at one stable fixed point on the manifold (left, colored dots). The ‘Set’ pulse sends trajectories beyond the saddle point, so that trajectories, after quickly converging to the manifold, evolve at different speeds towards the stable fixed point. Grey line: readout direction. **B** The projection of the different trajectories on the output direction generates a ramping signal with different speeds. The output time interval corresponds to the crossing of a fixed threshold. **C** Relation between cue amplitudes and output intervals in the designed model. Different cue amplitudes effectively modulate the output interval, as required by the task. **D** Trajectories of the designed solution for the MWG task. During the measurement and delay epochs (left, middle), the trajectories stay on the *κ*_1_−*κ*_2_ plane. The rank-three connectivity generates a sphere-like manifold. Within the ring of the manifold on the *κ*_1_ − *κ*_2_ plane, there are stable fixed points with very slow dynamics in their surroundings, that can be used for storing the signal. Left: during the measurement epoch, the first input initiates a slow transient trajectory. Middle: At the beginning of the delay, the trajectories corresponding to different input intervals decay to different states in the vicinity of the stable fixed point, and remain mostly unchanged during the rest of the delay period. Right: the ‘Set’ pulse sends trajectories towards the other side of the manifold. The trajectories evolve at different speeds depending on their location on the manifold during the delay period. **E** Readout activity for the different input intervals. A ramping signal with different slopes is produced after the ‘Set’ pulse. The output time interval corresponds to the crossing of a fixed threshold. **F** Relation between input interval and output interval for two different delays. The network produces different intervals based on the estimated input interval. Parameters are available in shared code.

**Supplementary Figure 13.**
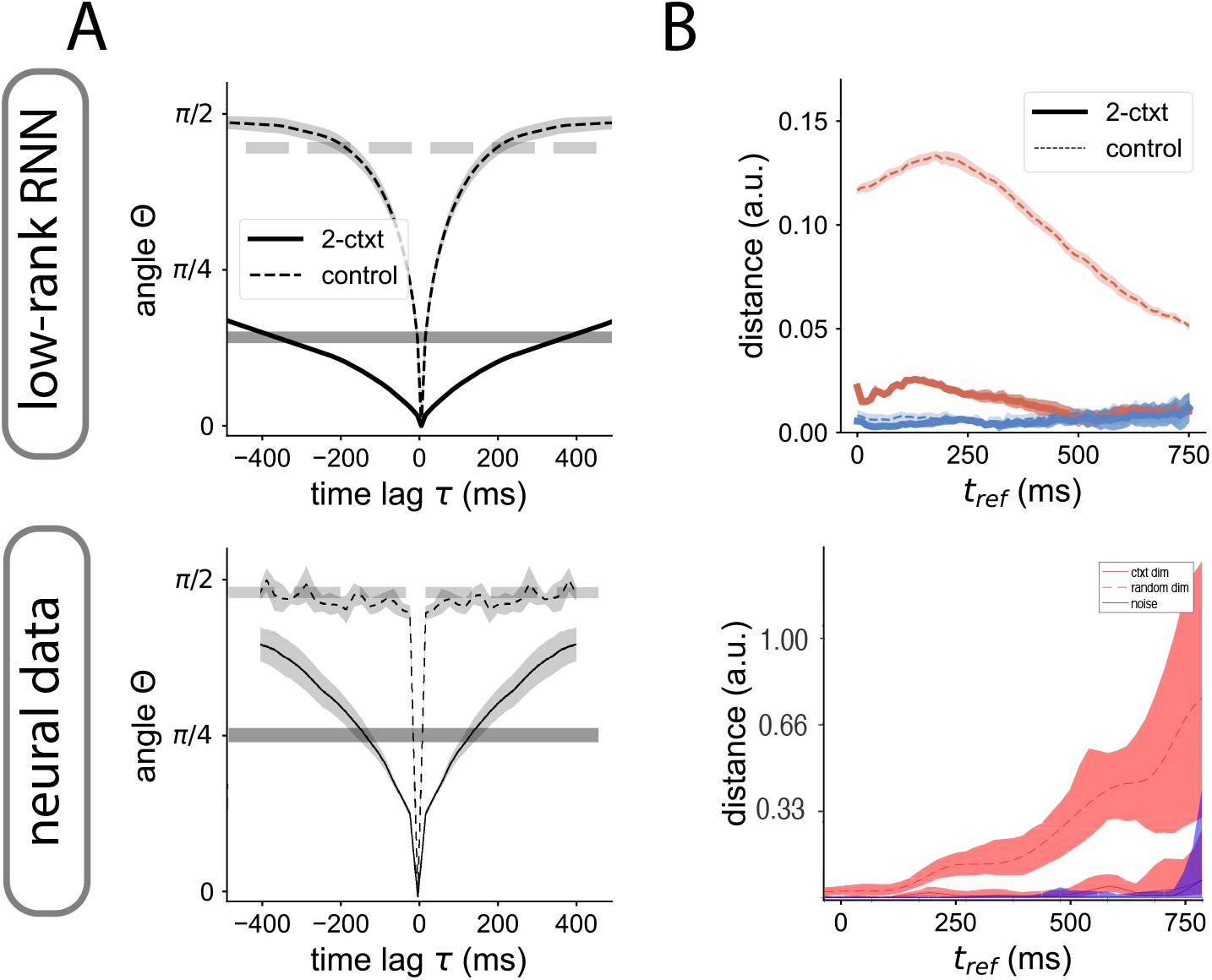
**A** Angle (in absolute value) as a function over time. Horizontal lines represent the mean value, shown in Fig. 6A. **B** Distance between trajectories as a function over time during measurement. The average value of the distance over time is shown in Fig. 6B.

